# Identifying Phenotype-Indicative Molecules from the Structure of Biochemical Reaction Networks

**DOI:** 10.1101/2025.10.21.683796

**Authors:** Yong-Jin Huang, Atsushi Mochizuki, Takashi Okada

## Abstract

Multistability in chemical reaction networks (CRNs) is a key mechanism underlying the diversity in cellular phenotypes. Identifying the steady state a cell occupies requires measuring molecular concentrations. This in turn raises a fundamental question: which chemical species should be selected to reliably distinguish among multiple steady states? We introduce a network-structural approach for identifying *indicator species*—a subset of species whose concentrations alone suffice to identify which steady state the system occupies. By decomposing a CRN into subnetworks based on structural criteria and applying topological degree theory, we show that the concentrations within certain subnetworks uniquely determine those of all remaining species. These subnetworks thus provide indicator species that distinguish all possible steady states. Crucially, our method relies only on stoichiometric and regulatory information, without requiring kinetic parameters. Implemented via a computational algorithm, the framework is validated through numerical simulations and applied to biochemical networks. This work provides a principled strategy for steady-state identification in high-dimensional biochemical networks from partial observations, with potential applications in single-cell data analysis and biomarker discovery.

## 1 Introduction

Cells exhibit remarkable phenotypic diversity, even if they share the identical genome. While RNA-seq data has traditionally been used to characterize phenotypes, the cellular phenotypic heterogeneity may not be fully determined by the transcriptional processes. Recent advances in single-cell technologies have shifted attention to metabolite profiling as a means of defining *metabolic phenotypes* [1, 2, 3, 4, 5, 6, 7]. Metabolite concentrations reflect the cumulative outcome of genetic regulation and cellular adaptation; the metabolic profiles offer a direct readout of phenotypic states [8, 9, 1, 10] and capture certain aspects of cellular heterogeneity that may be obscured in other omics, enabling deeper insights into cellular mechanisms.

A major challenge in single-cell metabolic phenotyping posed is the massive number of metabolites. Ideally, comprehensive quantification of all metabolites would enable precise differentiation of phenotypes (Fig. 1 (a)). However, such exhaustive analysis remains technically demanding due to the vast chemical complexity [8, 10]. High dimensionality poses a particular challenge, as it can bias multivariate analyses in metabolomics [11, 12]—commonly through the *curse of dimensionality* —and this challenge is further compounded by the typically limited number of cell samples available in practice [6]. This leads to a fundamental question: *which metabolites are most informative and sufficient to capture the system’s phenotypic diversity?* Addressing this question would greatly facilitate cellular phenotyping by focusing on a key subset of metabolites (Fig. 1 (b)). From a dynamical-systems perspective, the problem can be reformulated as identifying a subset of variables that fully characterizes multistability—the coexistence of multiple steady states that underlies the emergence of distinct phenotypic states under comparable conditions.

**Figure 1.**
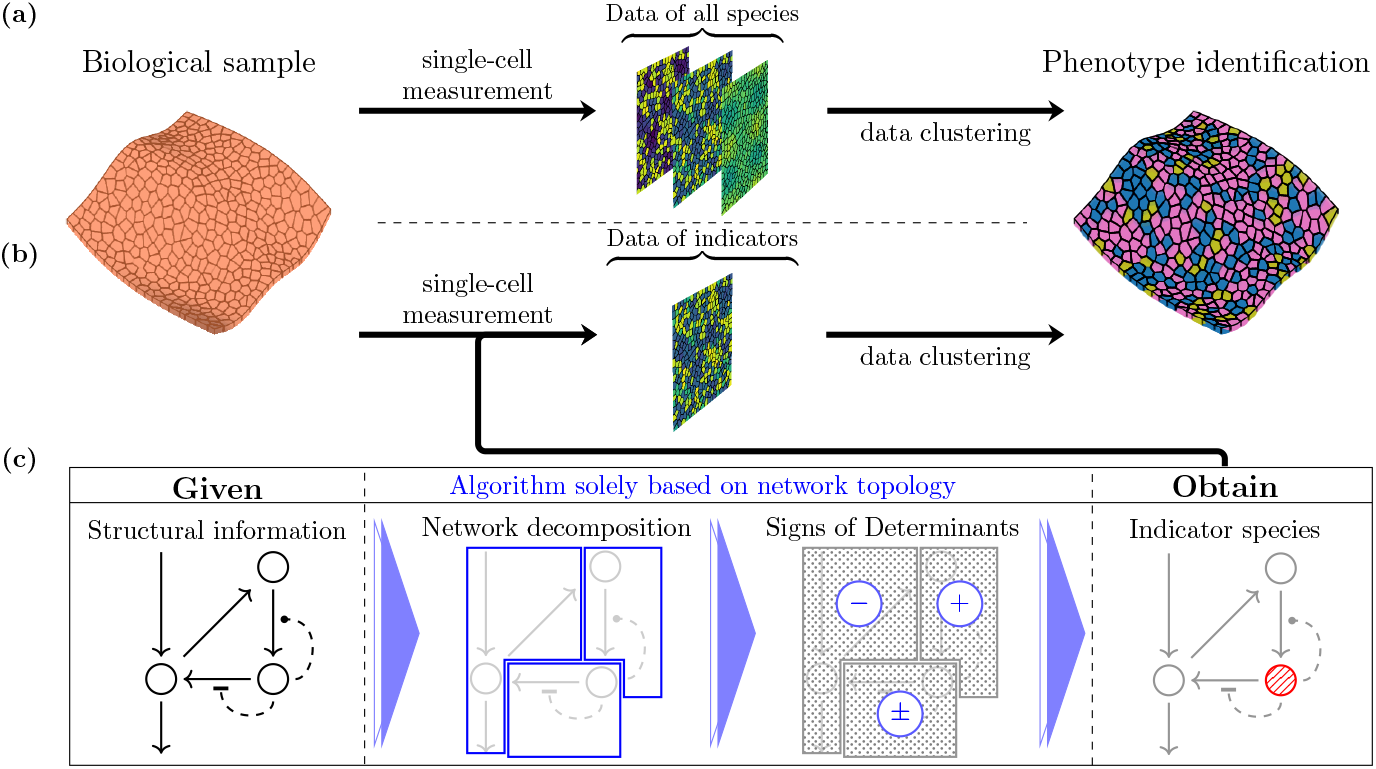
Distinguishing multistable phenotypes via indicator species. (a) Conventional pipeline, in which all available chemical species are measured and used for cellular phenotyping. (b) When indicator species are known, both measurement and classification tasks can be facilitated. (c) Our proposed theory and algorithm identify the indicator species directly from the network structure.

Multistability has long been studied in chemical reaction network (CRN) theory since the 1970s [13, 14], leading to a rich body of results that provide criteria for the existence of multiple equilibria [15, 16, 17]. Prominent among the criteria for precluding multistationarity are deficiency theory and the injectivity criterion, which form the bases for numerous methods that rely on network structure [18, 19, 20, 21, 22, 23, 24, 25, 26]. Other graph-theoretic approaches, though not specific to chemical systems, use feedback loop analysis based on Jacobian sign structure [27, 28]. However, these methods primarily address whether multistability can occur, rather than which species can serve as informative indicators of distinct steady states.

Several theoretical frameworks have been proposed to address the problem of identifying indicator variables. One class of approaches focuses on variable elimination, aiming to reduce model complexity by identifying species whose concentrations can be uniquely determined from others [29, 30, 31, 32]. In this context, the variables retained after elimination could be viewed as indicators. However, these approaches typically assume power-law kinetics, limiting their practical applicability to biochemical reaction networks. Another line of work, developed in the context of gene regulatory networks, uses feedback vertex sets (FVSs) to identify representative nodes that govern system dynamics [33, 34]. FVS-based approaches are purely structural and applicable to a wide range of dynamical systems. Nonetheless, they are not specifically designed to detect or rule out multistability in CRNs, as they do not fully leverage CRN-specific structural features relevant to steady-state analysis, such as stoichiometric interdependencies.

To address these gaps, we propose a general, structural method for identifying a subset of *indicator species* that reliably represents the system state and distinguishes phenotypic diversity arising from multistability (Fig. 1 (c)). In prior work, a Jacobian-equivalent matrix (see Eq. (5)) that explicitly encodes network topology was introduced; based solely on network structure, it was applied to analyze steady-state sensitivity [35, 36, 37, 38, 39], bifurcation phenomena [40, 41, 42], and network reduction [43]. The present work builds on this structural foundation by developing new theoretical results for a different problem: distinguishing among steady states. Unlike existing methods, our approach requires only stoichiometry and qualitative regulatory information and does not rely on kinetic details, aligning well with the practical demands of analyses based on biological pathway databases.

The remainder of this paper is organized as follows. The Results section begins by outlining the objective and introducing essential terminology. We then present our theoretical framework and corresponding algorithm, followed by numerical demonstrations that validate and evaluate the robustness of the approach. We conclude the section with an application to Reactome [44], an open-source pathway database, to illustrate practical utility. The Discussion section reflects on the broader implications and potential future directions. Detailed procedures for reproducing the results are provided in the Methods section.

## 2 Results

### 2.1 Discriminating System States: Terminology and Goal

Consider a chemical reaction network 𝒢 = (𝒳, ℛ) consisting of *M* chemical species, 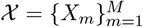, and *N* chemical reactions, 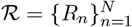

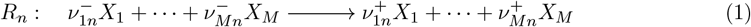

for *n* = 1, 2, …, *N*, where 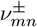’s stand for the coefficients in the chemical reactions. Defining the stoichiometric matrix as 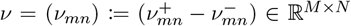 we describe the system’s dynamics via the ordinary differential equation

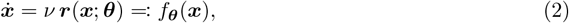

where 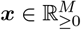 denotes the concentrations of all chemical species, 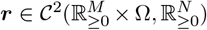 represents the reaction rates. The vector ***θ*** ∈ Ω ⊆ ℝ^*P*^ denotes *P* model parameters and is incorporated into the rate functions to account for biological variability; even among cells exposed to comparable conditions, slight heterogeneity in, e.g., temperature or enzyme activity can lead to variation in reaction kinetics. We assume that the domain Ω is assumed to be path-connected.

When the stoichiometric matrix *ν* is not full-ranked, it possesses nontrivial left-null vectors, indicating conservation relations among concentrations. For simplicity in presenting our results, the main text assumes *ν* is full-ranked, implying no such conservation constraints on the dynamics. Results on networks where *ν* is not full-ranked are provided in Appendix B.

The focus in this work is on multistability in the equilibrium dynamics, with equilibria, denoted by 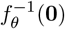, interpreted as phenotypic states of the system (e.g., the three clusters in Fig. 2 (c) and, for formal definition, Appendix A). To clarify our objective of identifying phenotype-indicating species, we introduce some notations and terminology. For a subset 𝒮 ⊆ ℕ_≤*M*_ := {1, 2, …, *M* }, define the projection

**Figure 2.**
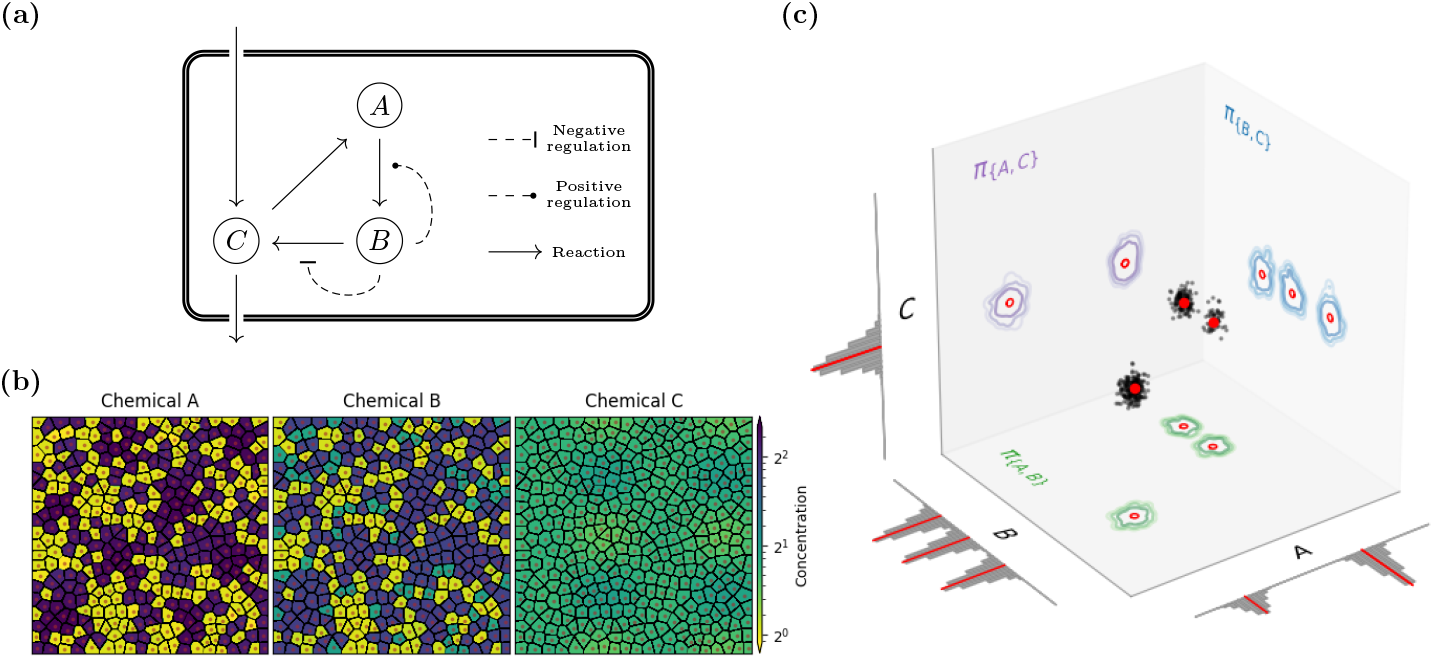
A three-species reaction network as an illustrative example. (a) The structure of the intracellular network. (b) Simulated tissue composed of cells, each assigned a parameter value in a position-dependent manner. The concentration of chemical C changes smoothly across the tissue. Chemical B reveals three distinct phenotypic states, while chemical A fails to distinguish two of them. (c) Scatter plot of cell states with contour plots of 2D projections to 2-species subsets and histograms of 1D projections to 1-species subsets. Red dots indicate three equilibrium states under a specific parameter setting, demonstrating that none of the subsets ∅, {*A*}, {*C*}, or {*A, C*} qualify as indicator subsets, as the projections cause overlaps among the equilibrium states.

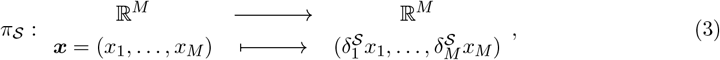

where for *m* ∈ ℕ_≤*M*_,

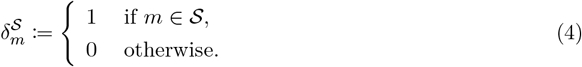

In conventional empirical approaches, determining the phenotypic state of a given steady state ***x***^*′*^ requires exhaustive measurements of the chemical concentrations; if concentration data for all species were available, one could identify which subset of chemical species is sufficient for determining the phenotype. However, this is impractical due to the difficulty of measuring every species. Furthermore, if the data are noisy, including irrelevant species may increase the risk of misidentification. Thus, our aim is to *a priori* find a subset 𝒮 ⊆ ℕ_≤*M*_ such that the phenotypic state can be determined solely by the values *x*^*′*^_*s*_ for *s* ∈ 𝒮.

#### Definition 1 (indicator subset)

*A subset* 𝒮 ⊆ ℕ_≤*M*_ *is an indicator subset if and only if*

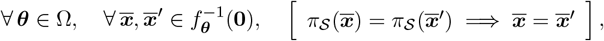

*that is, it is sufficient to determine the equilibrium points by chemicals in* 𝒮.

In the context of this work, one can equivalently (proved in Appendix) define indicator subsets by

#### Definition 2 (indicator subset)

*A subset* 𝒮 ⊆ ℕ_≤*M*_ *is an indicator subset if and only if, for any* ***θ*** ∈ Ω, *we always have* 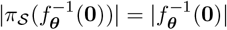; *namely, the equilibrium points stay distinguishable after the projection to* 𝒮.

For clarifying the term, Fig. 2 (a-c) gives a three-species example network, in which {*B*} and its supersets are indicator subsets, while ∅, {*A*}, {*C*}, {*A, C*} are not. Note that, for any system, the set of all chemical species is trivially an indicator subset.

The indicator subsets generally depend on both the stoichiometric matrix *ν* and the detailed kinetics (i.e., ***r***(***x***; ***θ***)). However, the latter is typically unknown for most intracellular reactions. Therefore, instead of assuming specific kinetics, we only assume that the signs of the partial derivatives 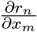’s are known. For instance, 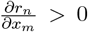 if *X*_*m*_ is a reactant of *R*_*n*_, and, in some cases, regulatory information is available (e.g., 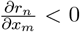 when *X*_*m*_ is an inhibitor of *R*_*n*_) from databases such as KEGG [45] and MetaCyc [46]. Thus, our goal is to find a non-trivial indicator subset 𝒮 solely based on the qualitative information (*ν* and signs of 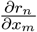) in a general kinetic setting.

### 2.2 Identification of Indicator Subsets: Theory and Algorithm

A structural approach to studying chemical reaction networks originates from the sensitivity analysis initially explored by Mochizuki and Fiedler [36, 35]. This approach has since expanded to bifurcation analyses [40, 41] and network reduction techniques [43]. The theory is centered on the use of a square matrix ***A*** constructed from network data. Specifically,

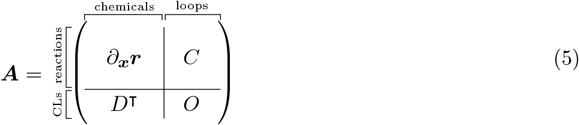

where the columns of *C* (resp. *D*) provide a basis for ker *ν* (resp. ker *ν*^⊺^), with its interpretation being the loops (resp. conservation laws) in the network system [38]. Indeed, in the main text, the lower blocks *D* and *O* should be absent due to the assumption that ker *ν*^⊺^ is trivial; nonetheless, the more general form is presented here to address cases with non-trivial ker *ν*^⊺^, as discussed in Appendix C.

It has been demonstrated in previous works how a chemical reaction network can be decomposed in the structural approach. According to the network topology, it is feasible to rearrange the rows and columns of the matrix ***A*** into a block upper triangular form [37, 38, 40, 39]; that is,

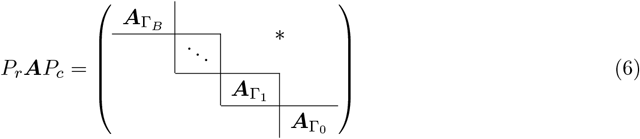

with 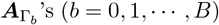 being square blocks, where *P*_*r*_ and *P*_*c*_ are permutation matrices. Since each chemical/reaction is associated with a column/row of the matrix, the block upper triangular form defines a network decomposition 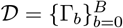 such that 𝒢 = ∪_Γ∈*D*_ Γ, where each Γ_*b*_ = (𝒳_*b*_, ℛ_*b*_) represents a subnetwork. If the column (resp. row) associating a chemical *X*_*m*_ (resp. reaction *R*_*n*_) lies on the square block 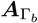, then we set *X*_*m*_ ∈ 𝒳_*b*_ (resp. *R*_*n*_ ∈ ℛ_*b*_). We name such a network decomposition as a *structural decomposition*. In addition, for the block upper triangular form as given in Eq. (6), we call it a *structural decomposition form* (of ***A*** with respect to 𝒟). This form does not have a unique canonical representation, since the labeling of Γ_*b*_’s is not unique, but this does not affect the final outcomes in our theory.

Our main theoretical result is that an indicator subset can be identified by means of structural decompositions. Specifically, we have

#### Theorem 1 (Main)

*Given a structural decomposition* 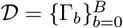 *such that, for any b*^*′*^ ∈ [0, *B*], *the subnetwork* 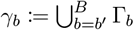 *is a buffering structure with no emergent conserved quantities. Put*

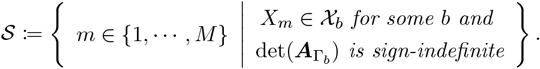

*Then*, 𝒮 *is an indicator subset of the chemical reaction network. Moreover, let* 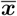 *represent the equilibria and put* 𝒮^*c*^ := {1, …, *M* }\𝒮, *then* 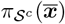 *is unique up to* 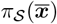 *and* ***θ***.

The proof is provided in the Appendices. The statement involves two terms not yet introduced: buffering structures [37, 38] without emergent conserved quantities [43] and sign-indefiniteness. Buffering structures without emergent conserved quantities are a special class of subnetworks, characterized by specific structural conditions, where the steady-state sensitivity to parameter changes within the subnetwork can be analyzed using only local structural information, independent of the outside network. To keep the main discussion accessible, we omit the formal definition here; see Appendix C for the details. The term sign-indefiniteness, as used in this paper, is defined by

#### Definition 3 (sign-indefiniteness)

*A mapping* 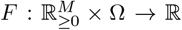 *is said to be sign-definite if the sign of F is constant over* 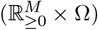, *namely*,

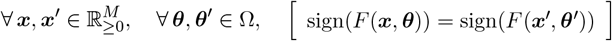

*Besides, F is said to be sign-indefinite if it is not sign-definite*.

In our theory, the sign-indefiniteness is judged for each det 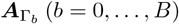, which is a polynomial of 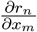 ‘s over ℝ. Using the information about the signs of derivatives 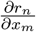 ‘s, it is feasible to identify whether each of det 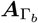’s is sign-definite for any kinetics. The main theorem shows that, for metabolite profiling, the metabolites for classification can be predetermined according to the network structure.

Based on this theorem, we developed an algorithm for identifying phenotype indicator species in chemical reaction networks. The core idea is to decompose the network into subnetworks Γ_*b*_’s and sequentially eliminate them in decreasing order of *b*, by using a reduction technique developed in Ref. [43], after verifying the sign-indefiniteness of det 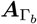’s. A complete set of pseudocodes is provided in Appendix D, and an implementation of the algorithm is available as a Python package `crnindsp` on GitHub (https://github.com/chromian/crnindsp.git).

### 2.3 Applications to a running example

To illustrate our novel method, consider a five-species chemical reaction network

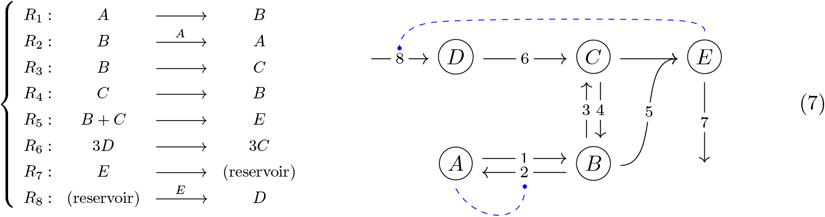

with the qualitative regulatory information given by *r*_2*A*_ *>* 0 and *r*_8*E*_ *>* 0, where 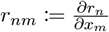. With the information above, the matrix ***A*** and its rearranged form are given by

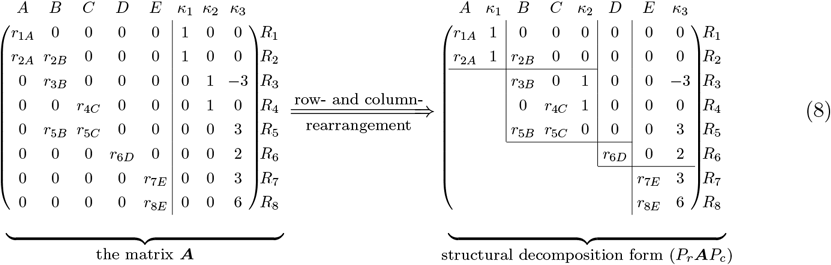

The rearranged matrix suggests a structural decomposition as Γ_0_ = {*E*}, Γ_1_ = {*D*}, Γ_2_ = {*B, C*}, and Γ_3_ = {*A*}. According to **Theorem 1**, the indicator subset consists of the chemical species corresponding to sign-indefinite diagonal blocks. The determinants det 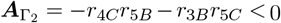 and det 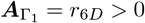 are both sign-definite, while det 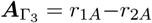 and det 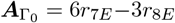 are sign-indefinite. Therefore, we conclude that 𝒮 = {*A, E*} is an indicator subset of the system.

To test our method’s prediction, we construct an ODE model of the chemical reaction network with specified forms of rate functions (see Sec. 4.1 in Methods for details). We simulate *N*_cell_ = 350 “cells,” each defined by randomly sampled parameters and initial conditions, yielding equilibrium concentrations 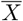. Observational data 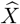 is generated by adding measurement noise to log-transformed 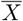. Hereafter, we refer to such a dataset 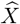, consisting of *N*_cell_ cells, as a tissue sample. The noise-free data 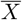 clearly separates into three clusters, interpreted as three phenotypic states (Fig. 5 in Appendix E). We use this clustering result as ground truth to evaluate how well subsets of chemical species in the noisy observational data 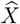 can correctly identify the underlying phenotypic states.

A defining property of an indicator subset, as formalized in Definition 1, is that it should enable clear separation of phenotypic states. To assess this property for each subset of chemical species, we project the observational data 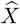 onto that subset and compute the overall silhouette score (the average of silhouette scores across all cells/points) relative to the ground-truth clustering; higher values indicate stronger within-cluster cohesion and greater between-cluster separation. Figure 3 shows the projected concentrations and the corresponding silhouette scores for a representative tissue sample. We perform this analysis for 100 independently-generated tissue samples, and the resulting distributions of silhouette scores are shown in Fig. 4 (a) (b). As illustrated in Fig. 3, projections onto arbitrary subsets of chemical species often lead to substantial overlap among phenotypic states, making accurate classification difficult. For instance, heuristics may suggest using {*B, C*} because of their high centrality in the network, but two of the three phenotypes exhibit substantial overlap in the {*B, C*} plane. In contrast, projection onto the indicator subset {*A, E*} yields a much clearer separation of the three phenotypes, as predicted.

**Figure 3.**
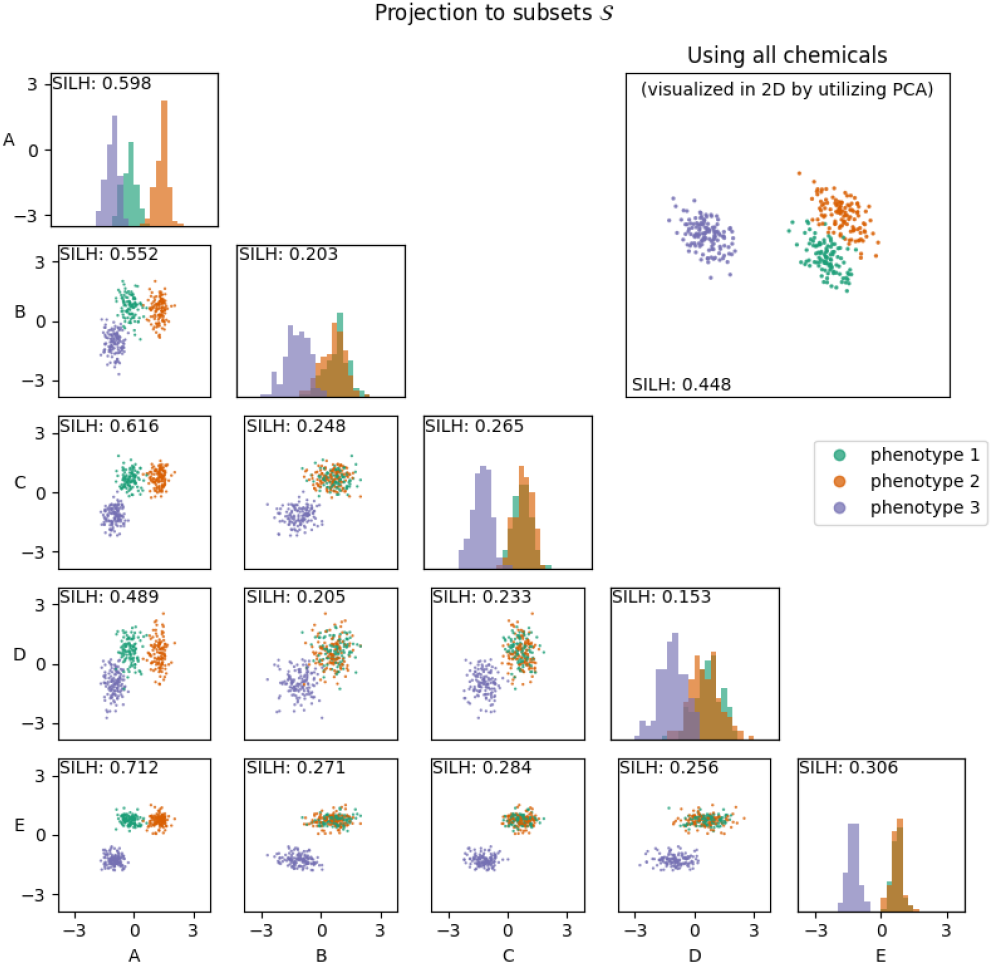
Projected observational data of chemical concentrations for a tissue sample of *N*_cell_ = 350 cells. Each point represents a cell, colored by the ground truth phenotype obtained from clustering noise-free data. The smaller panels show projections of the log-transformed data onto subsets of one or two chemical species, with silhouette scores (SILH) indicating phenotype separability. The larger panel displays the full five-species data, visualized using the first two principal components; the corresponding SILH is computed using all five variables.

**Figure 4.**
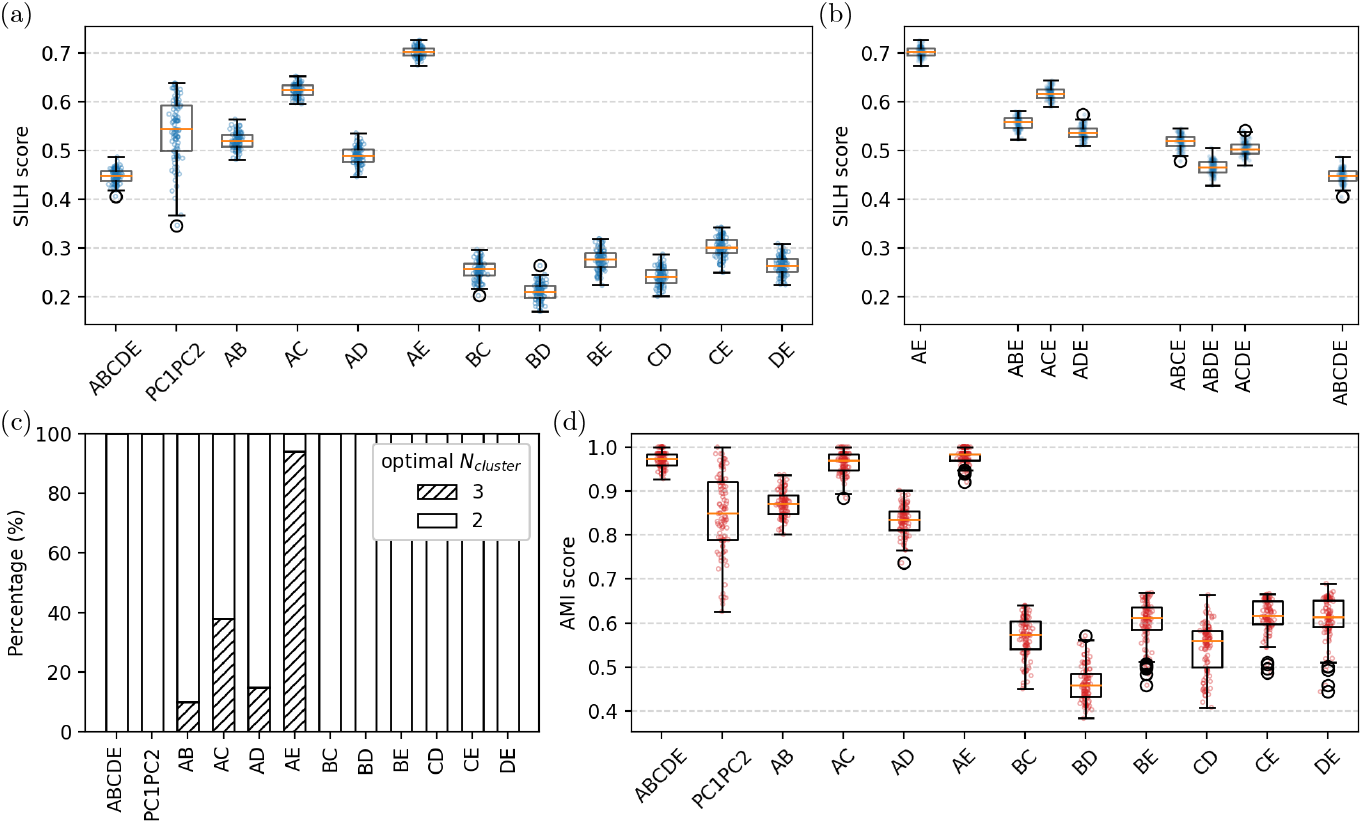
Validation of variables as indicators of phenotypic states using 100 replicated tissue samples. (a) Silhouette scores obtained under projections to the universal set (i.e., all chemicals), two principle components, and all the 2-species subsets. (b) Silhouette scores obtained under projections to the all the 𝒮 such that {*A, E*} ⊆ 𝒮. (c) The percentage of empirically determined clusters number, indicating how accurately each projection recovers the true number (that is, 3) of phenotypes. (b) Distributions of Adjusted Mutual Information (AMI) scores (given *N*_cluster_ = 3), quantifying the agreement between the ground truth and clustering results based on the projected observational data.

It is remarkable that the silhouette score for {*A, E*} significantly exceeds that of the full set {*A, B, C, D, E*} (Fig. 4 (a)), despite the latter also being an indicator subset. Projecting the five chemical species into two dimensions using PCA improves the silhouette score relative to the full set, but still performs worse than {*A, E*} (see PC1PC2 in Fig. 4 (a)). These suggest that, in the presence of noise, increasing the number of chemical species used may actually degrade classification performance. Supporting this, as we progressively enlarge subsets containing {*A, E*}, silhouette scores consistently decline (Fig. 4 (b)), illustrating the curse of dimensionality.

We next examine whether the observational data projected onto each subset can reveal the correct number of phenotypic clusters—corresponding to the criterion in Definition 2, an alternative formulation of an indicator subset. A commonly employed unsupervised approach to estimating the number of clusters *N*_cluster_ is to perform clustering (e.g., spectral clustering) for various values of *N*_cluster_ and select the one that yields the highest silhouette score (see Methods). Following this approach, we empirically determine *N*_cluster_ for each subset of chemical species by clustering the projected 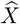 and selecting the value that maximizes the silhouette score. By performing this for 100 replicated tissue samples, we evaluate how frequently each value of *N*_cluster_ is selected (Fig. 4 (c)). The results show that projection onto the subset {*A, E*} most frequently identifies the correct number of clusters, *N*_cluster_ = 3. Specifically, under the parameter setting we used, it identifies the correct number of clusters in more than 90 out of 100 trials, while no other subset exceeds 40% accuracy.

Finally, we evaluate the accuracy of phenotyping for each subset by performing clustering on the projected observational data and comparing the clustering results with the ground truth. As the comparison metric, we use the Adjusted Mutual Information (AMI), which reaches 1 when all cells are classified consistently with the ground truth and approaches 0 for a random guess (see Methods). Because the AMI is sensitive to the number of clusters used—and this number is often misestimated for subsets other than {*A, E*}—we compare the AMI scores across the subsets by fixing the number of clusters to the correct value *N*_cluster_ = 3, rather than estimating it empirically. This ensures a fair and conservative comparison and avoids underestimating performance due to incorrect cluster numbers. Evaluating the AMI scores for 100 replicated tissue samples, we obtain the results shown in Fig. 4 (d). The AMI scores show that classifications according to {*A, B, C, D, E*}, {*A, C*} and {*A, E*} outperform all others. Moreover, Wilcoxon signed-rank tests show that {*A, E*} performs significantly better the other two subsets (Table 1).

**Table 1:** One-side Wilcoxon signed-rank tests for further testing the validation of {*A, E*} as an indicator subset in comparison to two other comparably good subsets illustrated in Fig. 4 (b).

| Alternative hypothesis | $p$-value of Wilcoxon signed-rank test |
| --- | --- |
| $\text{AMI}[\{A, E\}] > \text{AMI}[\{A, B, C, D, E\}]$ | $p \approx 1.4 \times 10^{-6}$ |
| $\text{AMI}[\{A, E\}] > \text{AMI}[\{A, C\}]$ | $p \approx 3.5 \times 10^{-13}$ |

With this small network for method illustration and verifications, we manually constructed the matrix ***A*** and derived the indicator subset based on the main theorem. In fact, our algorithm is a scalable and applicable to large biological networks in databases, and we demonstrate the application in the next subsection.

### 2.4 Applications

We implemented the algorithm to identify indicator species in biochemical reaction networks using Reactome, an open-source, peer-reviewed pathway database [44]. While the database does not provide kinetic information, it offers the stoichiometry and qualitative regulatory information, i.e., the structural information sufficient for our method. Reactome networks may contain both proteins and low-molecular-weight chemicals/metabolites. Because proteomic and metabolomic processes often occur on different timescales, we excluded proteins from our analysis (see Methods). Table 2 lists the analyzed networks along with the corresponding indicator species identified by our method. Although experimental validation of these predicted indicator species is still lacking, we present our results as a reference for colleagues, highlighting which species may best represent the overall state of each system.

**Table 2:**
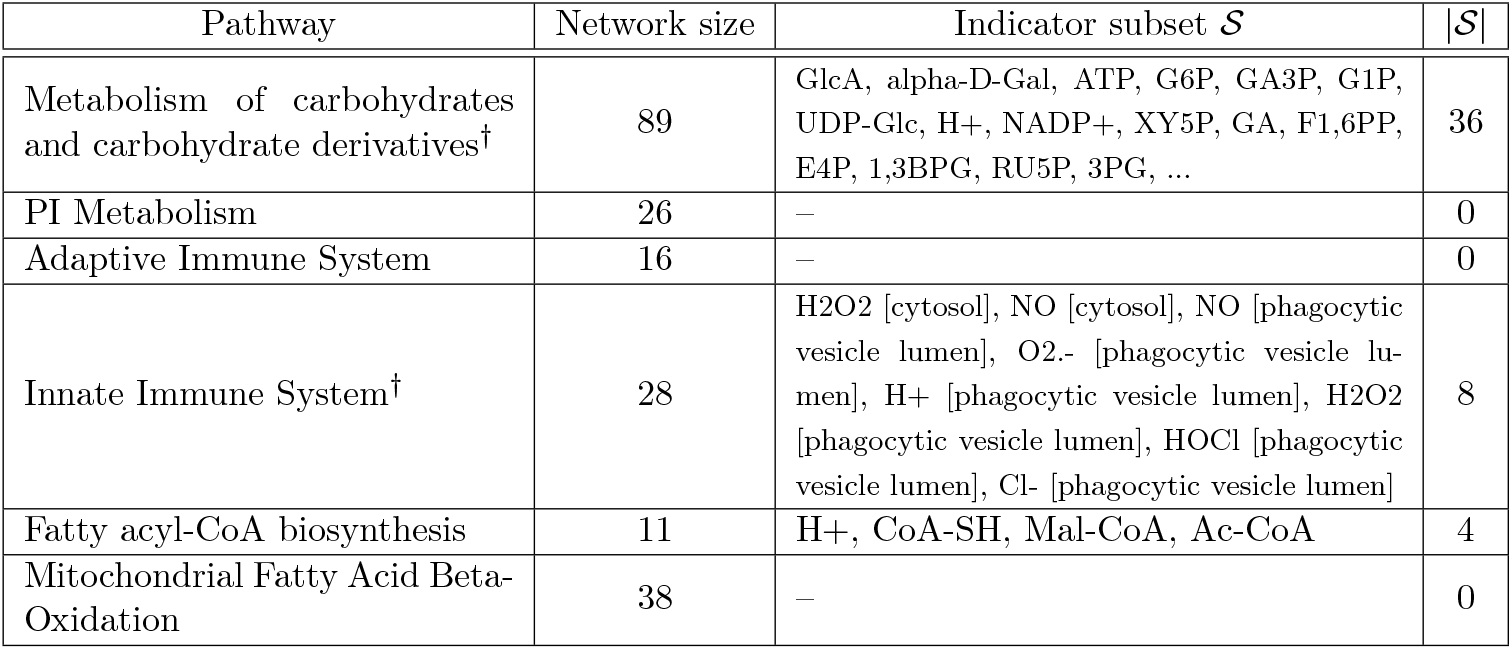
The indicator subsets determined by the proposing method. The listed networks are arbitrarily selected from Reactome for *Homo sapiens*. Networks with structural singularities are indicated by †.

Several networks extracted from the database were *structurally singular*, which is defined as the condition det(***A***) ≡ 0 for any ∂_***x***_***r*** of which sign pattern agrees with the given regulatory information. Such a condition is not expected to occur, as it implies the absence of any (asymptotically stable) steady states in the system (By Corollary 1 in [42]). Therefore, the occurrence of a singularity suggests that the network information is incomplete, potentially due to missing regulatory information or reactions. The singularity of a network can be contributed to some singular subnetwork(s) (that is, Γ_*b*_ with det 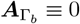), which are the origin of the misinformation.

A network’s singularity can result from the presence of one or more singular subnetworks (denoted Γ_*b*_), for which det 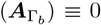. These subnetworks are presumed to be the origin of the inconsistency or misinformation in the database. For a given subnetwork Γ_*b*_ exhibiting det 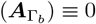 within the erroneous presumed network, it remains unclear whether the determinant is sign-definite (which implies that the chemicals in Γ_*b*_ may be negligible) or sign-indefinite. To ensure cautious interpretation in practical applications, we include the chemicals associated with such singular subnetworks in the indicator subset 𝒮.

Our results (Table 2) show that, in real-world biochemical networks, not all chemical data are required for metabolic phenotyping. Both measurement (i.e., quantifying chemical levels) and classification can be facilitated by our method, which substantially reduces the number of chemicals to be assayed, as suggested by Table 2. Interestingly, some of pathways yield empty indicator subsets, which means that the network dynamics do not exhibit multistability, and cells with the same gene expression levels must possess the same metabolic profile.

## 3 Discussion

In this study, we addressed an underexplored problem in CRN: identifying variables informative of multistability. Unlike prior approaches [16, 21, 23, 24] focused on establishing conditions for multistability, our method targets state distinguishability, independent of specific kinetic assumptions. Building on topological degree theory and structural analysis, we established a rigorous framework in which such indicator sets can be identified and provided an efficient algorithm for their computation. Numerical validations and applications to curated biochemical networks demonstrated the practical relevance and computational tractability of our framework. While our motivation stemmed from single-cell metabolic phenotyping, our results are broadly applicable across systems governed by reaction network dynamics, offering a general tool for identifying steady states of chemical reaction networks.

We emphasize the strengths of our method in identifying a subset of chemical species sufficient for analyzing the equilibrium dynamics of chemical reaction networks. The stoichiometric structure offers a distinct advantage: it provides clear and well-defined structural information—an asset often unavailable in other natural systems—and enables the reduction of system complexity to a lower-dimensional representation. Our effort to leverage such a structural feature facilitates analysis and reduces the burden of data collection. From a broader perspective, our method can be interpreted as a form of dimension reduction tailored to chemical systems. This is particularly valuable, as high-dimensional data analysis is often challenged by noise and the so-called curse of dimensionality. In the numerical investigation in this present paper, for instance, clustering results are significantly more accurate when using the identified indicator subset rather than the full set of species. An additional strength of our framework is the interpretability of the dimension reduction. Unlike conventional techniques such as PCA or UMAP, which often obscure physical meaning through projection into abstract feature spaces, our method preserves the physical identity of species. Thus, our reduced representation retains mechanistic interpretability while achieving substantial simplification.

Despite these practical strengths, several limitations warrant discussion. First, our theory is intended to distinguish state diversity arising from multistability, rather than diversity along a single solution branch caused by parameter variation. In other words, to apply our method to real biological systems, cells should be exposed to similar environmental conditions. However, even under shared environmental conditions, extrinsic variability—such as differences in gene-expression levels—may still arise, thereby obscuring the intrinsic diversity reflected by the predicted indicator species. To safeguard against this, we recommend first identifying the cell types using transcriptomic and/or proteomic data. Our method can then be applied within each type to classify subtypes, thereby capturing the intrinsic diversity generated by the system’s dynamics (namely, multistability).

Second, when applying our indicator species framework to real-world pathway databases such as Reactome, we identified potential errors in network structures that may affect their reliability. Our previous work [42] shows that the determinant of the corresponding matrix ***A*** indicates the system stability. In our analysis (Table 2), we found det ***A*** ≡ 0 for certain networks, suggesting an unstable system, which is biologically implausible for cellular processes. This likely indicates incomplete network data, such as missing reactions or regulatory interactions. In our kinetics-free framework, singularity of ***A*** typically stems from zero rows or columns, which must correspond to a singular block 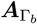 in the structural decomposition. This insight enables our framework to detect structural deficiencies, offering a tool for refining pathway databases and guiding experimental validation.

Finally, our current framework does not extend to oscillatory dynamics. Because it relies on equilibrium dependencies among chemical species, we provide no theoretical guarantees for systems exhibiting limit-cycle behavior. While empirical evidence for oscillations in metabolic networks is scarce, our method’s limitation in handling them suggests a potential diagnostic: If the theoretically predicted indicators fail to produce clear phenotypic separation, this may indicate that cells are following oscillatory rather than fixed-point trajectories.

A key direction for advancing the present theory lies in exploring whether the indicator subset can be further minimized. Specifically, we hypothesize that it suffices to select one representative species from each of sign-indefinite subnetworks in the structural decomposition. This idea is grounded in the center-manifold theorem, which suggests that the equilibrium manifold’s dimension, determined by the right-nullity of the matrix ***A***, reflects the minimal variables needed. An analogous but rigorous argument shall hold for subnetworks, implying that a single species per sign-indefinite subnetwork could represent its dynamics.

Inspired by the challenges of single-cell metabolomics, we envision our method as a transformative tool for experimentalists and data scientists tackling cellular phenotype classification. By identifying an indicator subset of species that capture CRN steady states, our method simplifies high-dimensional data analysis while preserving mechanistic interpretability. We invite researchers to apply this framework to single-cell datasets and make effective use of pathway databases, which remain underutilized despite the considerable effort invested in their creation. Its kinetics-free approach offers a powerful, accessible tool for advancing systems biology and related fields.

## 4 Methods

### 4.1 The data generation for the 5-species chemical reaction network

For the verification of our theory, we constructed an ODE model for the example network as given in Eq. (7). Specifically, for the dynamics given by

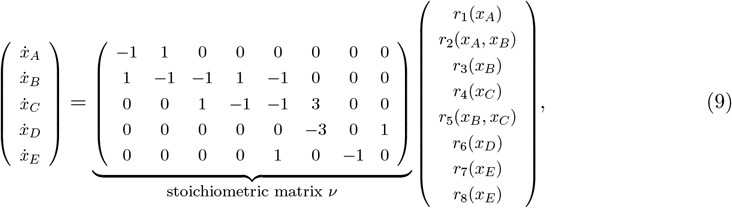

and we considered

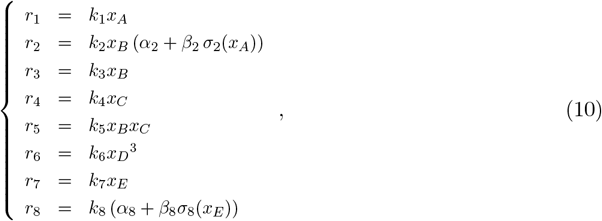

where *σ*_2_ and *σ*_8_ are sigmoid functions, and *k*_·_, *α*_·_, and *β*_·_ are positive constants. A power-law kinetics was basically taken as the strategy of modeling, whereas sigmoid functions were considered for the regulation *r*_2*A*_ and *r*_8*E*_, according to the network information. We put *α*_2_ = 0.65, *α*_8_ = 0.343, *β*_2_ = 3.25, *β*_8_ = (5 − 0.343). The sigmoid functions realizing the regulatory *r*_2*A*_ and *r*_8*E*_ are determined by

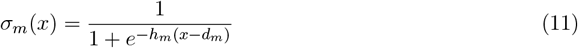

for *m* = 2, 8, where *h*_2_ = 18, *h*_8_ = 15, *d*_2_ = 1.55, *d*_8_ = 2.

With this model, we generated *N*_cell_ = 350 data points in 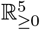 as the initial states of biochemical reaction networks for *N*_cell_ cells in a tissue. The model was run for each cell for a while long enough for the relaxation to equilibria. To better simulate real-world conditions, we accounted for parameter variations among cells in the following numerical experiments. Specifically, for each reaction *j* (1 ≤ *j* ≤ 8) in each cell *c* (1 ≤ *c* ≤ *N*_cell_), let *K*_*jc*_ denote the random variable for the parameter *k*_*j*_ in the *c*-th cell. Similarly, for each chemical *X*_*m*_ in each cell *c*, let *X*_*mc*_ represent the random variable for the initial concentration of the chemical in that cell. We assume 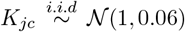 and 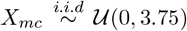, where 𝒩 and 𝒰 denote the Gaussian and uniform distributions, respectively. By numerically solving the ODEs for all cells and allowing the system to relax to equilibrium points, we obtain a data matrix 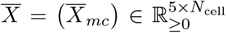, where each column vector represents the system state of a cell. Under these conditions, three equilibrium states are observed (Fig. 5 in Appendix E), with each cell converging to one of them. However, due to parameter variations, the state variable values at the same equilibrium state differ slightly among cells. To further align the model with real-world scenarios, we introduce measurement errors into the observation process. Specifically, we define 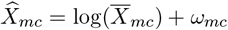, where 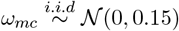. This results in a data matrix 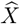, which represents the observed system states with measurement noise. In Appendix E, we provide additional demonstrations under other parameter settings.

### 4.2 Clustering and scoring

In the numerical demonstrations, we employ spectral clustering from the Python module `scikit-lear` as an unsupervised method for phenotype classification [47]. Broadly speaking, this method applies classical *k*-means clustering to distances derived from the spectral embedding of the input data (cf. [48]).

The silhouette score is adopted as the method for quantifying the internal consistency of data clustering. Given a set of data points {*y*_*i*_}_*i*∈ℐ_ and a partition (i.e., classification) {*C*_*l*_}_*l*∈ℒ_ of the dataset into clusters, the (individual) silhouette score for each point provides a measure of how well it fits within its assigned cluster compared to other clusters. For a data point *y*_*i*_ ∈ *C*_*l*(*i*)_, the following quantities are defined:

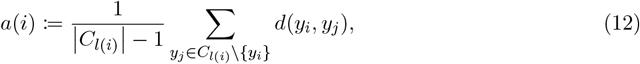

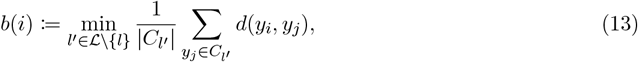

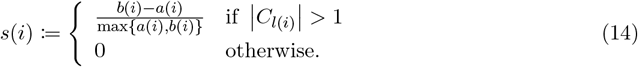

Here, *a*(*i*) represents the average dissimilarity between *y*_*i*_ and all other points within its assigned cluster, while *b*(*i*) denotes the lowest average dissimilarity between *y*_*i*_ and all points in any other cluster. The silhouette index *s*(*i*) ∈ [−1, 1] thus quantifies how dissimilar the data point *y*_*i*_ is from points in the nearest neighboring cluster, relative to the cohesion within its own cluster. Higher values of *s*(*i*) indicate better clustering quality for that point. Then, for a given classification {*C*_*l*_}_*l*∈ℒ_, the (overall) silhouette score is defined as the average of the individual silhouette scores computed over all data points in the dataset.

Clustering methods typically require the number of clusters to be specified in advance. A common unsupervised approach for determining this number is to select the value that maximizes the overall silhouette score. In the present study, the term empirically driven number of clusters refers to the selected value of |ℒ| that yields the highest overall silhouette score among the classification results {*C*_*l*_}_*l*∈ℒ_ produced by spectral clustering.

To evaluate classification performance based on different choices of observed target species, we adopt a widely used scoring metric called Adjusted Mutual Information (AMI), which quantifies the similarity between the ground truth and the predicted classification. Suppose that there are *N*_*c*_ many data points. Both 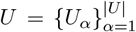 and 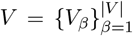 are partitions of the data points. Consider

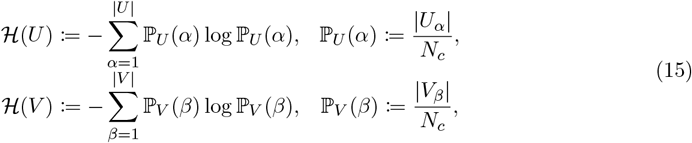

and the mutual information (MI) between *U* and *V* is given by

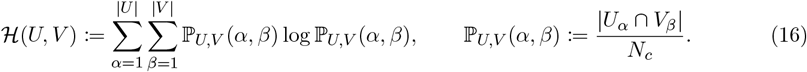

The Adjusted Mutual Information (AMI) adjusts MI to account for chance overlap between clusterings and is defined as

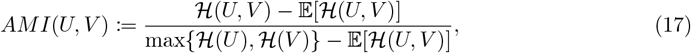

where the expectation 𝔼 [*MI*(*U, V*)] is computed under a hypergeometric model of randomness [49]. Assuming *U* represents the ground truth and *V* is the clustering result being evaluated, an AMI score of 1 indicates perfect agreement between *V* and *U*, while a score close to 0 suggests the clustering result is comparable to random guessing.

### 4.3 Data preprocessing for the implement on Reactome

To implement the proposed method on Reactome networks, we follow a specific preprocessing protocol tailored to the characteristics of the data and the theoretical assumptions of our framework. Reactome provides multi-omics network data, which comprise not only chemical species but also proteins and RNA/DNA. Since our method is derived within the framework of chemical reaction network theory, RNA/DNA species should certainly be excluded from the analysis.

Furthermore, due to substantial differences in the timescales of the governing dynamics of proteins and small molecules, we recommend that proteins and chemicals not be analyzed together within a single network. Mixing species with such disparate kinetic regimes can lead to violations of key assumptions, particularly the assumption of dissipativity, which is essential for our theorems (see Appendix A). Accordingly, users are advised to apply the method to protein-only and chemical-only networks separately. In our demonstration, we restrict attention to chemical species by selecting only those annotated with the Systems Biology Ontology (SBO) term SBO:0000247 (simple chemical).

Each network analyzed is treated as a subnetwork extracted from a larger Reactome pathway. Our primary interest lies in investigating the potential for multistability among intermediate species within these subnetworks. To this end, we exclude boundary species, namely, species with no fluxes or no outfluxes. Both types are omitted from the analysis to focus on the internal dynamics of the system.

## Acknowledgements

We are grateful to Carsten Conradi, Hiroshi Kokubu, Yuhei Yamauchi, and Atsuki Hishida for helpful discussions and comments. This research was supported by the CREST program (grant no. JPMJCR1922, JPMJCR24Q4) of the Japan Science and Technology Agency (JST) (http://www.jst.go.jp/EN/index.html), Grant-in-Aid for Scientific Research on Innovative Areas (Grant No. 19H05670), Joint Usage/Research Center program of Institute for Life and Medical Sciences Kyoto University. T.O. was supported by JSPS KAKENHI (Grant No. JP22K03453, JP22K06347, 25K09723, 25K07168) and the RIKEN iTHEMS Program.

## A Appendix: Jacobian, Dissipativity, and Phenotypic states of Chemical Reaction Networks

Adopting the notations in the main text, we consider the dynamical system following the ODE

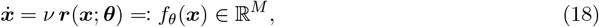

where *θ* ∈ Ω and Ω is a path-connected subset of ℝ^*P*^.

Let *D* ∈ ℝ^*M ×L*^ be a matrix such that its column vectors form a basis of ker *ν*^⊺^ ≃ coker *ν* = (ℝ^*M*^*/* im *ν*), where *L* = dim(ker *ν*^⊺^). In the main text, for simplicity, we consider the case in which ⟨*D*⟩ = ker *ν*^⊺^ = ∅. Nonetheless, this assumption does not hold in general, and for every system state ***x*** ∈ ℝ^*M*^, the *L* elements of the vector ***η*** := *D*^⊺^***x*** ∈ ℝ^*L*^ give all the conserved quantities, which are named so because

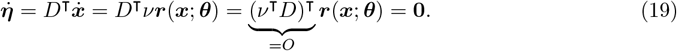

The values of the conserved quantities are determined by the initial conditions of the dynamics. Since they remain constant over time, they can be regarded as part of the external factors, incorporating ***η*** into ***θ*** like ***θ*** = (***η***, …) ∈ ℝ^*L*^ × ℝ^*P* −*L*^.

Since the existence of conserved quantities implies interdependencies among the variables ***x***, we may wish to extract independent dynamical degrees of freedom from ***x***. For this purpose, let *V* ∈ ℝ^*M ×M* −*L*^ be a matrix such that its column vectors form a basis of im *ν*. Consider the pseudo-inverses *V* ^*†*^ and *D*^*†*^ which are given by

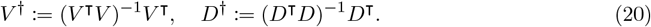

Then, columns of (*V* ^*†*^)^⊺^ and (*D*^*†*^)^⊺^ are linear independent, and one can further see that ⟨(*D*^*†*^)^⊺^⟩ = ⟨*V* ⟩^⊥^ = (im *ν*)^⊥^ (∵ *D*^*†*^*V* = *O* and dim ⟨(*D*^*†*^)^⊺^⟩ = *M* − dim ⟨(*D*^*†*^)^⊺^⟩). Analogously, ⟨(*V* ^*†*^)^⊺^⟩ = (ker *ν*^⊺^)^⊥^. The basis matrices of the linear subspaces considered so far give us an linear isomorphism

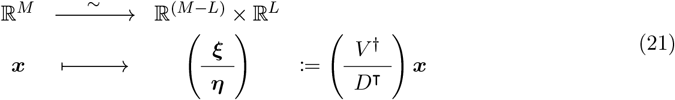

and its inverse is given by ***x*** = *V* ***ξ*** + (*D*^*†*^)^⊺^***η***. With such a change of coordinate, one sees that the system dynamics is sufficiently described by

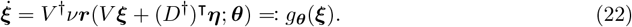

Note that the function *g*_***θ***_ is well-defined without being provided with ***η*** as an argument, because ***η*** is specified by ***θ***.

Since the system’s dynamical behaviors are reflected only by ***ξ***, when talking of the Jacobian matrix, we are considering

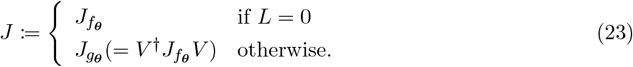

Note that when *L* = 0, for the arguments all above, we can put *V* = *I*; consequently, we have *f*_***θ***_ = *g*_***θ***_ and ***x*** = ***ξ***.

Such a modification of the Jacobian is to circumvent the problem that *L* eigenvalues always vanish due to the presence of conserved quantities even when the system is at a stable state. Since 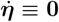, the system dynamics only relies on the dynamics of ***ξ***, and hence the stability of a system state should be determined by the modified Jacobian instead when *L*≠ 0.

### Definition 4 (Critical and regular values)

*Consider a mapping* 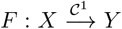. *A point y* ∈ *Y is said to be a regular value of F if and only if, for any x* ∈ *F* ^−1^(*y*) ⊆ *X, we have* det *J*_*F*_ |_*x*_≠ 0. *A point y* ∈ *Y is a critical value if and only if it is not a regular value*.

In this work, the interest is the equilibria of the chemical reaction network, so a following question is whether **0** is a regular value for the considered dynamics. In the practical situation of metabolic phenotyping, the data points represent stable equilibria; namely, for any ***x*** such that *f*_***θ***_(***x***) = **0**, the Jacobian is non-singular. This characterizes the parameter region that should be considered, and we thus assume that **0** is a regular value for ***θ*** almost everywhere in Ω. Note that ***η*** = *D*^⊺^***x*** represents the conserved quantities, which is time-invariant and treated as parameters, so the regularity is defined by *g*_***θ***_, not *f*_***θ***_. In summary, for the practical situation, we have the assumption

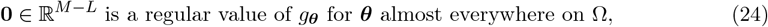

in which, by “almost everywhere”, we mean that the collection of parameters unsatisfying the property is of Lebesgue measure zero.

The notions above are introduced for (1) formally defining the phenotypic states of chemical reaction networks and (2) proving that the collection of equilibria is a finite set.

### A.1 Phenotypic states

Consider

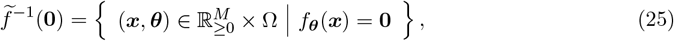

which is the collection of pairs of fixed points and its corresponding parameter(s).

#### Definition 5 (Phenotypic states)

*Phenotypic states are defined to be the equivalence classes of the equivalence relation* ≃ *given as follow: for* 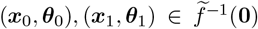, *we write* (***x***_0_, ***θ***_0_) ≃ (***x***_***1***_, ***θ***_***1***_) *if and only if there exists a continuously differentiable path* 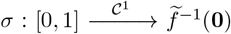 *such that*

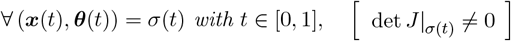

*and σ*(0) = (***x***_0_, ***θ***_0_), *σ*(1) = (***x***_1_, ***θ***_1_).

That is, if there is a continuously differential path on the equilibrium manifold from a point to another point in the state-parameter space such that *J* is not singular, then we say they are of the same phenotype of the chemical reaction network.

One may wonder whether this definition of phenotypic states aligns with those identified through clustering analysis of equilibria, which is the practical situation we are considering. Recall that ***θ*** ∈ Ω is a parameter vector of a CRN in cells in a tissue, and Ω ⊆ ℝ^*P*^ is assumed to be path-connected. Since ***θ*** represents the micro-environment conditions, the parameter variation δ***θ*** is supposed to be small for a cell and a neighbor of it. By our definition, the two cells are of the same phenotypic states only if the state difference δ***x*** → **0** when we smoothly let δ***θ*** → **0**. In such a procedure of parameter adjustment, the Jacobian *J* is always non-singular, hence the implicit function theorem shows that a phenotypic state can be expressed a continuously differential parameterization 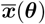 of system state by the parameters. This thus guarantees that each of the phenotypic states, denoted by 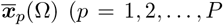, where the finiteness is shown in the next subsection), are path-connected and that they are mutually disjoint when Ω is small enough. In conclusion, the parameter variation (more formally, the diameter of Ω) is assumed to be small for cells and their neighbors in the practical case, clustering according to ***x*** does agree with our definition.

### A.2 Dissipativity and the finiteness of equilibria

Other than defining the singularity of system states and the phenotypic states, the linear isomorphism is also used for the formal definition of *dissipativity* of a chemical reaction network. We give the definition according to [50, 51], but using our own notations for self-consistency.

#### Definition 6 (Dissipativity)

*A considered chemical reaction network is said to be dissipative if and only if, for any* ***θ*** ∈ Ω, *there exists a compact set* 𝒦_***θ***_ ⊆ ℝ^*M*^ *such that, for any trajectory* ***x***(*t*) *with* ***x***_0_ = ***x***(0), *we have d*(*ω*(***x***_0_), 𝒦_***θ***_) = 0, *where* 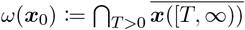, *and d*(·, ·) *is the Hausdorff distance*.

This assumption applies broadly to biological systems as it effectively prevents biologically unrealistic scenarios of unbounded growth. In this study, we put a stronger but still biologically reasonable assumption:

#### Definition 7 (Radial dissipativity)

*A considered chemical reaction network is said to be radially dissipative if and only if*,

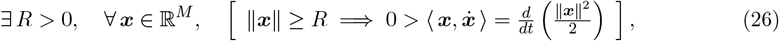

*where* ⟨ ·, · ⟩ *denotes the inner product of* ℝ^*M*^ *and* ∥·∥ *is the norm*.

This assumption is a sufficient condition of the dissipativity, since one can see that ∩ = *B*_*R*_(**0**) (the ball of radius *R* centered at **0**) is the desired compact global forward-invariant set. Notice that ∥***x***∥ can be taken as a measure for the total abundance of chemical concentrations, ensuring that 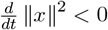 asymptotically prevents its unbounded growth.

A philosophical question may follow as, considering the fact that a biochemical network is usually a subnetwork of a larger network (e.g., a pathway), what kind of (sub-)networks are appropriate to be assumed dissipative. We apply the assumption to any biochemical network that *behaves as a complete system* in the long term; that is, if any system-specific parameter (e.g., enzyme activity) undergoes perturbations, and it eventually does not affect the concentrations of chemicals out of the system, then we assume it to be radially dissipative. In Appendix C, we will specify the idea stated in this paragraph more concretely.

#### Theorem 2

*The radially dissipative condition (26) of* ***x*** *is satisfied only if* ***ξ*** := *V* ^*†*^***x*** *also satisfies the condition, that is*,

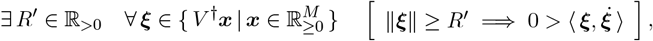

(proof) Let *R* ≥ 0 be the positive number as in (26). Arbitrarily choose a 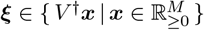 such that ∥***ξ***∥ *> R*. By the fact that *D*^*†*^*D* = *I, V* ^*†*^*V* = *I*, and *D*^*†*^*V* = *O*, for 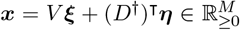, we have ∥***x***∥^2^ = ***x***^⊺^***x*** = ***ξ***^⊺^***ξ*** + ***η***^⊺^***η*** ≥ ***ξ***^⊺^***ξ*** = ∥***ξ***∥^2^ ≥ *R*^2^. Then, (26) is followed by 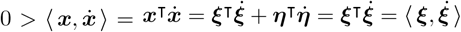, since 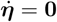.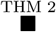

#### Theorem 3

*Consider the dynamics* 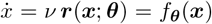 *of a chemical reaction network that is dissipative. When* **0** *is a regular value of g*_***θ***_, *then* 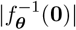 *is finite*.

(proof) For a given ***θ***, the vector ***η*** of conserved quantities is also fixed if *L* := dim(ker *ν*^⊺^)≠ 0. For 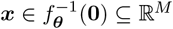, the linear isomorphism (***ξ, η***) I→ ***x*** given by ***x*** = *V* ***ξ*** + (*D*^*†*^)^⊺^***η*** implies that there is a 1-1 correspondence between 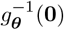 and 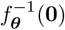. On the other hand, if *L* = 0, then we simply take *V* = *I* for the isomorphism ***x*** = *V* ***ξ***. In summary, it is sufficient to show 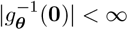. Since 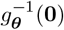 is a preimage of a continuous map *g*_***θ***_ of a closed set {**0**}, one sees that 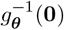 is also closed. By the radial dissipativity and Theorem 2, 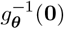 is in a compact set, thus being bounded. Then, with Heine–Borel theorem, 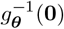 is seen to be compact. Since **0** is a regular value of *g*_***θ***_, the inverse function theorem tells us that 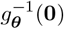 is discrete. Therefore, if 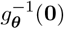 is not finite, we can write 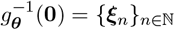, and by the inverse function theorem, there exists a sequence {*V*_*n*_}_*n*∈ℕ_ of open sets such that 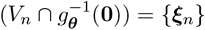 for every *n* ∈ ℕ. Apparently, {*V*_*n*_}_*n*∈ℕ_ forms an open cover of the compact set 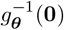, and hence there exists a finite subsequence 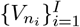 of {*V*_*n*_}_*n*∈ℕ_ such that 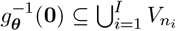, so we have 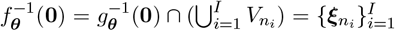, which leads to a contradiction to the assumption that 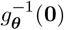 is an infinite set. Hence, 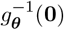 is finite. 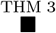

#### Theorem 4

*Let* 𝒮 ⊆ ℕ_≤*M*_. *When* 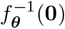 *is a finite set, the following statements are equivalent:*

1. 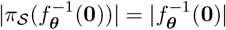
2. 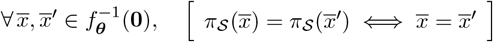

(proof) Suppose that *2*. holds, then there exists an bijection between 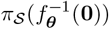 and 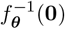, and thus *1*. holds. Now we suppose that *2*. does not hold, then *π*_𝒮_ is surjective but not injective, and by the finiteness of 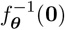, we conclude 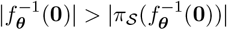, that is, the statement *1*. is false.The above concludes that the statements *1*. and *2*. are equivalent. 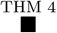

## B Appendix: Multistability analysis rooted in the Degree theory

Before continuing the discussion on

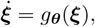

to give a slightly more general introduction to the degree theory, we instead consider

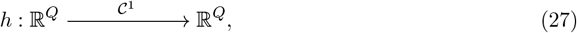

where 𝒪 ⊆ ℝ^*Q*^ is open and bounded. When this is the case, we can define

### Definition 8 (Brouwer degree)

*For an open and bounded set* 𝒪 ⊆ ℝ^*Q*^ *and a regular value* ***y*** ∉ *h*(∂𝒪), *where* ∂𝒪 *denotes the boundary of* 𝒪, *the Brouwer degree d*_*B*_(*h*, 𝒪, ***y***) *is given by*

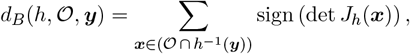

*where the sum over an empty set is defined to be zero. The shorthand notation d*_*B*_(*h*, 𝒪) := *d*_*B*_(*h*, 𝒪, **0**) *is used for* ***y*** = **0**.

This is not the original definition of Brouwer degree but comes from a proposition stating that, for a regular value, the value of the degree can be obtain as a finite sum of signs of Jacobian determinants [52]. The interest of this work is the case when ***y*** = **0**, which is a regular value of *h* = *g*_***θ***_, and we hence simply take it as a definition.

Brouwer degree serves as a powerful tool for multistability analysis since it can count the fixed points of dynamics. In particular, in this study, we make use the fundamental properties as follows:

[P0] If *d*_*B*_(*h*, 𝒪, ***y***) = 0, then 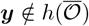 namely, when the degree is zero, there is no solution ***x*** in the closure of 𝒪 for ***y*** = *h*(***x***).

[P1] (**Normalization**) If *id* _*O*_ is the identity map on 𝒪 and ***y*** ∈ *h*(𝒪), then *d*_*B*_(*id* _*O*_, 𝒪, ***y***) = 1.

[P2] (**Homotopy invariance**) Let 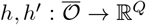 be two homotopy equivalent C^0^ functions via a continuous homotopy 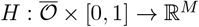 such that *H*(***x***, 0) = *h*(***x***) and *H*(***x***, 1) = *h*^*′*^(***x***), then ***y*** ∉ *H*(∂𝒪 × [0, 1]) =⇒ *d*_*B*_(*h*, 𝒪, ***y***) = *d*_*B*_(*h*^*′*^, 𝒪, ***y***).

In this regard, whether the Brouwer degree *d*_*B*_(*h*, 𝒪) takes the value of zero tell us whether or not there is a fixed point in 𝒪; moreover, if *f* is homotopic to the identity map, then there is a fixed point only when *d*_*B*_(*h*, 𝒪) = 1.

### Theorem 5

*If the system is radially dissipative, and the Jacobian determinant (*det *J) is signdefinite, then* 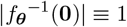 *for* ***θ*** *almost everywhere on* Ω.

(proof) Fix a ***θ*** ∈ Ω such that **0** is a regular value of *g*_***θ***_ (and hence, (−*g*_***θ***_)). By the radial dissipativity and Theorem 2, there exists a number *R >* 0 such that

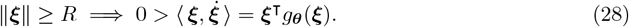

This gives 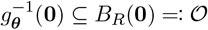. Let *id* be an identity map on 𝒪 and consider a homotopy

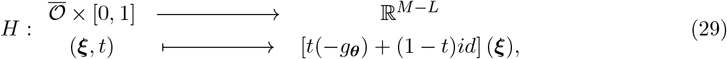

Apparently, *H* is a continuous homotopy between (−*g*_***θ***_) and *id*. Next, let ***ξ*** ∈ ∂𝒪 be arbitrarily given. Then, when *t* = 0, we surely have *H*(***ξ***, 0) = ***ξ***≠ **0** since ∥***ξ***∥ = *R >* 0. On the other hand, for any *t* ∈ (0, 1], if there exists any ***ξ*** ∈ ∂𝒪 such that **0** = *H*(***ξ***, *t*), then by the construction of *H* we see 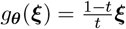, which leads to a contradition

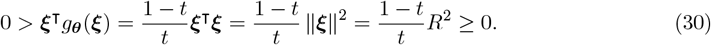

The above concludes that **0** ∉ *H*(∂*B*_*R*_(**0**) × [0, 1]). By normalization and homotopy invariance of the degree, we have

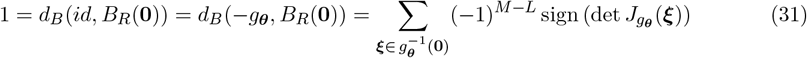

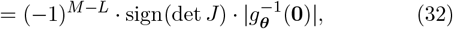

since det *J*_*g****θ***_ = det *J* is sign-indefinite by assumption. This shows that sign(det *J*) ≡ ( ^−^1)^*M*−*L*^ and that 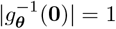. Since such kind of ***θ*** is almost everywhere on Ω, the proof is complete. 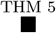

We are inspired by a pioneering work [50] to have come up with Theorem 5. The authors and their collaborators have a rich series of works on the multistationarity/multistability of chemical reaction networks. We acknowledge that not only the authors of the cited paper [50] and the collaborators but also many other colleagues greatly contribute to the theory of chemical reaction networks on multistability. Whereas, the proof above is given by following the one given in the supplementary material of [50]. In their derivations, the modification of Jacobian is different from ours, and the dissipative condition is relaxed. Readers carefully follow these technical (and maybe frustrating) parts are genuinely encouraged to check those pioneering works. Other than [50], one may like to start from, for instance, [19, 26].

These pioneering works focus on the question whether multistability is allowed by a parameter region, kinetics, or network structure. Meanwhile, our aim is to find the subset of variables to determine the state, where the network structure is given, and the other information is unclear. Therefore, in the next section, we are going to show how the theorem can be applied to subnetworks.

## C Appendix: Structural analyses using network topology

### The matrix *A*, Jacobian matrix, and buffering structures

Recall the matrix ***A*** suggested in the main text

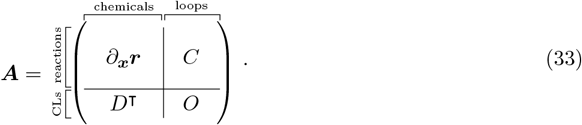

It is noteworthy that this matrix is suggested to study the sensitivity and stability problems of chemical reaction networks [35, 38, 41]. The matrix reveals the local behaviors of the dynamics because the matrix is associated with the Jacobian matrix, and a transforming formula between the two matrices is explicitly given in our previous work [42]. More specifically, we have the relation

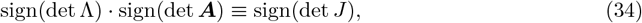

where Λ is a invertible constant matrix determined by *ν*, given by

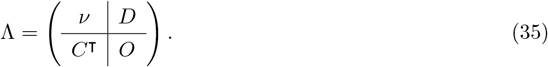

Based on this, rather than using det *J*, we can alternatively use det ***A*** for the definitions and theorems in previous sections.

The advantage of using the matrix ***A*** instead of the Jacobian matrix is the structural insights given by the structural approach. The matrix is known for the ability of identifying *buffering structures*, the subnetworks that can localize bifurcaiton behaviors and system responses to parameter perturbations [37, 40, 41]. A buffering structure is defined to be a subnetwork *γ* = (𝒳_*γ*_, ℛ_*γ*_) consisting of a set 𝒳_*γ*_ of chemicals and a set ℛ_*γ*_ of reactions such that it satisfies

1. (output-completeness) Reactions regulated by any chemical in 𝒳_*γ*_ are included in ℛ_*γ*_.
2. 0 = *χ*(*γ*) := |ℛ_*γ*_| − |𝒳_*γ*_| + (# conserved quantities involving *γ*) − (# loops in *γ*).

The statement here could appear ambiguous, since the last two terms in the definition of *χ*(*γ*) are not clearly defined math objects at this point. We do have a formal definition for the index *χ*, whereas it requires more notations for the specification, so we put the rigorous definition in the next subsection after introducing necessitated notations. Other than this graphical definition, there is actually an equivalent way of defining buffering structures:

#### Definition 9 (Buffering structures)

*Let γ be a subgraph of* 𝒢, *with γ*^*c*^ *denoting the complement of γ. If* {*γ, γ*^*c*^} *is a structural decomposition such that, for some permutation matrices P*_*r*_ *and P*_*c*_, *the structural decomposition form can be expressed as*

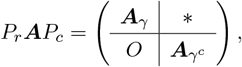

*then γ is called a buffering structure*.

Although the former definition is more frequently used in other papers of this structural analyses (e.g., [37, 40, 39, 41, 43]), the equivalence has been proven in a previous work [42]. Such a substructure is called a buffering structure because, when parameter perturbations are given within a buffering structure, in the long-term behavior of the equilibrium dynamics, the perturbation does not influence the rest of the system, but the system responses are confined within the substructure [37, 40]. This property is called the *law of localization* arising from buffering structures. In this regard, a buffering structure is an open subsystem unilaterally influenced by the rest of the system, and it behaves as a complete system in the long term. Therefore, with the argument in Appendix A, it is reasonable to assume that every buffering structure forms a system with dissipation when the rest of the system is considered as the external factors. A formal statement for this assumption is given later around Eq. (59), for this purpose we would need to prepare some notations first.

#### C.2 Structural reduction of chemical reaction network

A network reduction can be conducted by the eliminations of buffering structures [43]. For the sake of argument, we first consider a subnetwork *γ* = (𝒳_*γ*_, ℛ_*γ*_), which is not necessarily a buffering structure, and its complement is denoted by *γ*^*c*^ = (𝒳_*c*_, ℛ_*c*_). Let 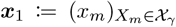 and 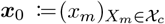 denotes the concentration levels of chemical in *γ* and *γ*^*c*^, respectively. Similarly, let ***r***_1_ := ***r***_1_(***x***; ***θ***) and ***r***_0_ := ***r***_0_(***x***; ***θ***) denote the rate functions of reactions in ℛ_*γ*_ and ℛ_*c*_, respectively. The ODE is thus rewritten as

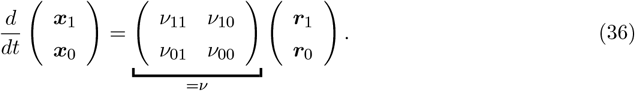

Let

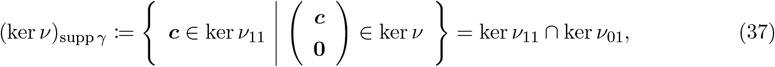

and we get well-prepared to give the graphical definition of buffering structures in a formal manner.

##### Definition 10 (Buffering structures)

*Adopt the notations above. The considered subnetwork γ is a buffering structure if and only if the following statements are satisfied:*

1. *(output-completeness)* 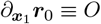.
2. *(zero index)* 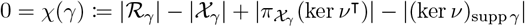

*in which | · | denotes the cardinality for a finite set, whereas for a linear subspace it denotes the dimension*.

Revisit Eq. (36). When the system is at an equilibrium point, we surely have 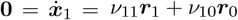, and thus

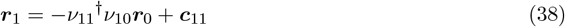

for some ***c***_11_ ∈ ker *ν*_11_. We first <u>**claim:** *γ* is a buffering structure with no emergent conserved quantity =⇒ ker *ν*_11_ ⊆ ker *ν*_01_,</u> leaving the proof of it and the definition of emergent conserved quantities later. With the claim, when *γ* is such a buffering structure, by substituting Eq. (38) into the second equation in the system Eq. (36), one obtains

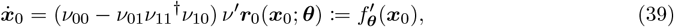

where the term *ν*^*′*^ := *ν*_00_ − *ν*_01_*ν*_11_^*†*^*ν*_10_ is the Schur complement (*ν/ν*_11_) of the block *ν*_11_ of the matrix *ν*, taken as the stoichiometric matrix of the reduced system. Besides, by the outputcompleteness, we see 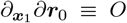, which means that we can write ***r***_0_ = ***r***_0_(***x***_0_; ***θ***) since it is independent of concentration levels of chemicals in the buffering structure. Besides, remark that for any 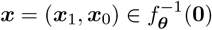, it must be satisfied that 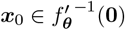; namely,

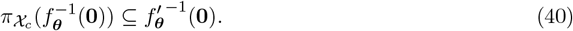

In verbal terms, the structural reduction ensures that any equilibrium of the original system, when projected onto the reduced state space, remains an equilibrium of the reduced system. This property of structural reduction will be used later.

Now we explain and prove the aforementioned **claim**. Notice that by applying rank–nullity theorem to the linear mapping 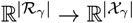 defined by *ν*_11_, one sees

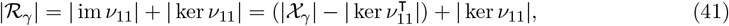

since ker 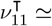 coker 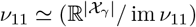. Hence, the index can be rewritten as

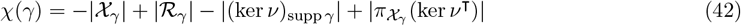

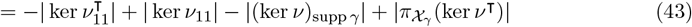

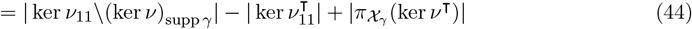

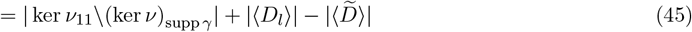

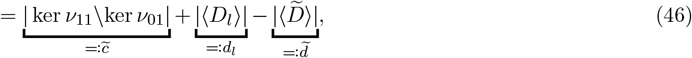

where

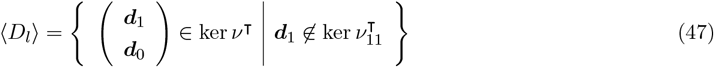

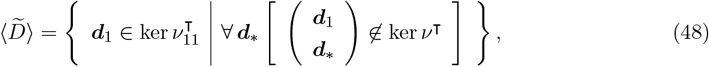

in which *D*_*l*_ and 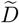 denote the matrix of which column vectors form bases of their spanning spaces, respectively. Note that 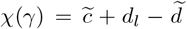, where the three numbers are non-negative integers. Specifically, the 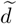 column vectors of 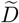 specify the *emergent conservation laws*, which appear to be conservation laws within the buffering structure but are actually not really conservation laws in the system. By saying the buffering structure has no emergent conserved quantity, we mean 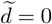, which is immediately followed by 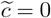, showing the **claim**.

##### Definition 11

*A buffering structure is said to be self-contained if it is a buffering structure with no emergent conserved quantities; namely*, 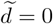.

In the present work, when talking of structural reduction, we mean a structural reduction with respect to a self-contained buffering structure.

##### Theorem 6

*If the considered chemical reaction network satisfies the dissipative condition (26), then a structurally reduced system following the dynamics (39) also satisfies the dissipative condition*.

(proof) By assumption, there exists some *R >* 0 such that for any ***x*** of which norm is larger than *R*, we have ⟨ ***x***, *f*_***θ***_(***x***) ⟩ *<* 0. For the reduced system, let 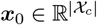 be an arbitrarily given vector such that ∥***x***_0_∥ *> R*. Then, putting ***x***_1_ := −(*ν*_01_*ν*_11_^*†*^)^⊺^***x***_0_, we apparently have ∥***x***∥ *> R* for ***x*** = (***x***_1_, ***x***_0_), *ν* ***r*** and hence

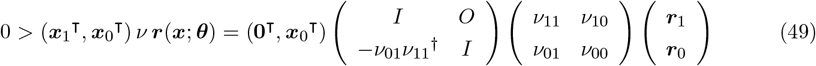

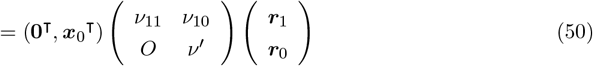

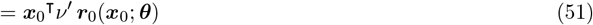

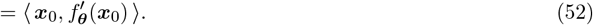

This completes the proof. 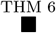

Before continuing to the next theorem for the theory of this work, we introduce a fact

##### Lemma 1

*For any system state* ***x***, *which is not necessarily an equilibrium, we have*

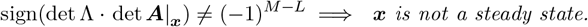

This lemma is proved in a previous work [42] and hence we skip the proof. We shall stay aware of the fact that we are deriving theorems for (bio-)chemical reaction networks in living systems, of which equilibrium points stand for phenotypic states. Among the equilibrium points, there must exist some steady one; otherwise, there exists no stable phenotypic state, and such kind of network should not have appeared as a considered network. In summary, there must exist at least a steady state, and sign(det Λ · det ***A***|_***x***_) = (−1)^*M*−*L*^ at the point. This is followed by a fact that det ***A***≢ 0, which we will use in the proof of the following theorem:

##### Theorem 7

*Let γ be a buffering structure, and say we have*

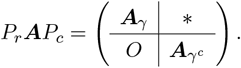

*Then, the* ***A****-matrix of the reduced system can be rearranged to be* ***A***_*γ*_*c by row- and column permutations*.

(proof) Since a structural decomposition form does not have a unique canonical representation but is flexible to row- and column-rearrangement, without loss of generality, we may write

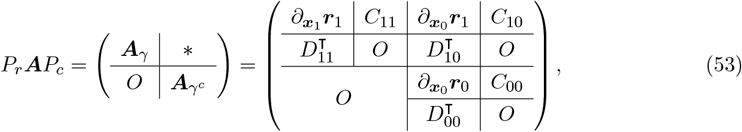

where

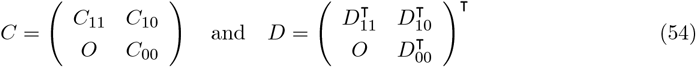

are matrices of which the column vectors respectively form a basis of ker *ν* and a basis of ker *ν*^⊺^. To prove the theorem, it is sufficient to show (1) the column vectors of *C*_00_ form a basis of ker *ν*^*′*^ and (2) the column vectors of *D*_00_ form a basis of ker *ν*^*′*⊺^. We first check whether the column vectors of *C*_00_ and those of *D*_00_ are both linear independent. Had it been not, then det ***A***_*γ*_*c* ≡ 0 due to the linear dependency, implying det ***A*** ≡ 0; nonetheless, as mentioned right before the statement of this theorem, det ***A***≢ 0. This concludes the linear independency of column vectors of both *C*_00_ and *D*_00_. Besides, note that we have

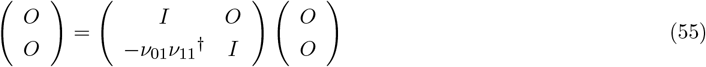

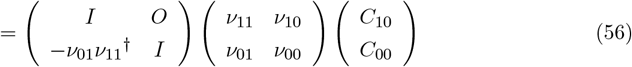

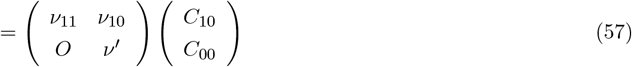

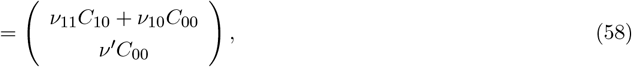

which means that *ν*^*′*^*C*_00_ = *O*, so the subspace ⟨*C*_00_⟩ spanned by columns of *C*_00_ is a subset of ker *ν*^*′*^. Check the dimension, then we see ⟨*C*_00_⟩ = ker *ν*^*′*^, according to the linear independency of column vectors. In summary, column vectors of *C*_00_ form a basis of ker *ν*^*′*^. One can analogously see that column vectors of *D*_00_ form a basis of ker *ν*^*′*⊺^. Now we have the proof. 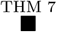

#### C.3 The derivation of the Main Theorem

By the identity given in Eq. (34), we can rewrite Theorem 5 into

##### Theorem 8

*If the system is radially dissipative, and* det ***A*** *is sign-definite, then* |*f*_***θ***_^−1^(**0**)| ≡ 1 *for* ***θ*** *almost everywhere on* Ω.

This is a special case that none of the chemicals in the considered network exhibit the multistability behavior, which means that we do not need the data (concentration levels) of any of them to distinguish the phenotypic states. That is, 𝒮 = ∅ is an indicator subset. What we are going to do is applying this theorem to *γ* and *γc*, so as to see whether they are responsible for the multistability behaviors; if not, then they can be excluded from 𝒮.

To do it, we first once again stress that the structural reduction keeps the concentration levels of remained chemicals at equilibria (Eq. (40)). Therefore, if the reduced system has a unique equilibrium point, then for any two equilibrium points 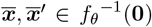, we must have 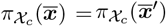, which implies that chemicals in *γc* are not responsible for the multistability behaviors, and we can exclude them from the consideration of indicator subsets. Besides, the radial dissipativity is adopted from the original system to the reduced system via a structural reduction (Theorem 6). Furthermore, a structural reduction also preserves the matrix ***A***; more specifically, the matrix associated with the reduced system 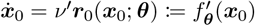 is exactly ***A***_*γ*_*c* (Theorem 7).

On the other hand, recall that we argued that a buffering structure forms a dissipative subsystem when considering the rest of the system is treated as its parameter. Specifically, we mean that, when fixing the outside concentrations 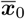 and ***θ***, the dynamical system of the buffering structure,

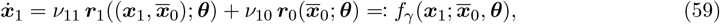

is assumed to satisfy the radial dissipativity.

##### Theorem 9

*Given a self-contained buffering structure γ and let its complement denoted by γc*. *If* det ***A***_*γ*_ *is sign-definite, then γ in general has a unique equilibrium up to* 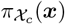 *and* ***θ***; *that is*, 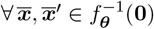, *we have* 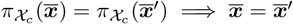 *for* ***θ*** *almost everywhere on* Ω.

(proof) Consider a system

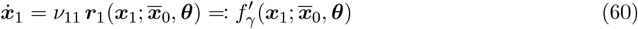

which shares the same Jacobian matrix 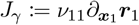 with the system given by Eq. (59). We can consider Eq. (60) as the dynamics of a chemical reaction network, where *ν*_11_ is the stoichiometric matrix. For such a chemical reaction network, we would like to construct its ***A***-matrix. More specifically, we <u>**claim**</u> that

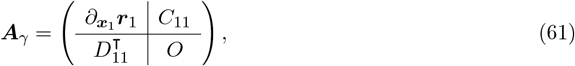

which is given in Eq. (53), is the ***A***-matrix for Eq. (60). To show the **claim**, it is sufficient to show that column vectors of *C*_11_ and *D*_11_ form bases of ker *ν*_11_ and ker 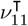. Since 0≢ det ***A*** = det ***A***_*γ*_ · det ***A***_*γ*_*c*, the column vectors of *C*_11_ and *D*_11_ are both linear independent. Note that *ν*_11_ is immediately followed by *ν*_11_*C*_11_ = *O*, and then the dimensionality shows that *C*_11_ does provide a basis for ker *ν*_11_. On the other hand,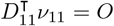; otherwise, there exits some ***d*** = (***d***_1_, ***d***_0_) ∈ ker *ν*^⊺^ such that ***d***_1_∉ ker 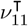, which means the dimension *d*_*l*_ of ⟨*D*_*l*_⟩ as given in Eq. (47) is nonzero, which is contradictory to the proved fact that 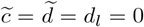 when *γ* is a self-contained buffering structure. The <u>**claim**</u> above is now proved. Next, put

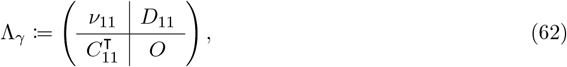

of which determinant can be seen to take a nonzero value. Then the identity Eq. (34) gives sign(det Λ_*γ*_) · sign(det ***A***_*γ*_) ≡ sign(det *J*_*γ*_). By assumption, det ***A***_*γ*_ is sign-definite, and hence so is det *J*_*γ*_. Applying Theorem 5, we see that the system given by Eq. (59) has a unique equilibrium for ***θ*** almost everywhere on Ω. With Theorem 3 and Theorem 4, the proof is compete.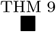

As stated in the main theorem, suppose that we are given a structural decomposition 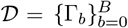 (where Γ_*b*_ = (𝒳_*b*_, ℛ_*b*_)), 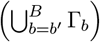 is a self-contained buffering structure for any *b*^*′*^, and that the structural decomposition form is given as in Eq. (6).

##### Theorem 10

*Suppose that* det 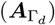 *is sign-definite for some* (𝒳_*d*_, ℛ_*d*_) = Γ_*d*_ ∈ 𝒟. *Given an indicator subset* 𝒮 *such that* ∪_*d>b*≥0_ 𝒳_*b*_ ⊆ 𝒮, *it follows that* (𝒮\𝒳_*d*_) *is also an indicator subset*.

(proof) Let ***θ*** ∈ Ω be arbitrarily given. Put

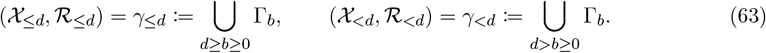

Notice that (*γ*_≤*d*_)*c* = ∪_*B*≥*b>d*_ Γ_*b*_ is either an empty set or a self-contained buffering structure, so one can apply the structural reduction by eliminating the *γ*. By the preservation of a structural reduction, for the reduced system we have the structural decomposition form

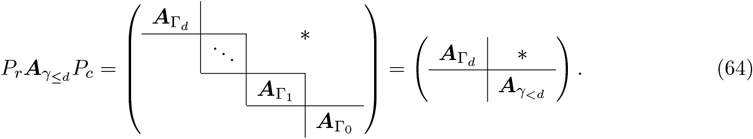

This shows that, in the reduced system, Γ_*d*_ is a self-contained buffering structure, with *γ*_*<d*_ being its complement. In addition, recall that the reduced system is radially dissipative. Moreover, by assumption, det 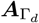 being sign-definite. In summary, we can apply Theorem 9 to the reduced system and hence conclude

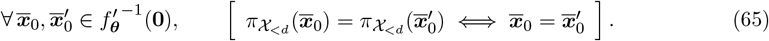

By the property Eq. (40), we further see that the same thing happens in the original system; more specifically, for any 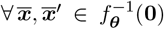, we certainly have 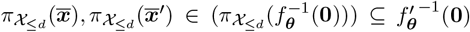, and thus

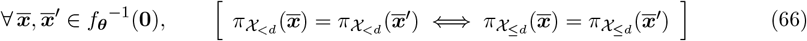

Now, suppose that 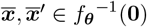 are two points such that 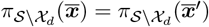, which means

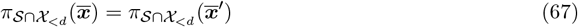

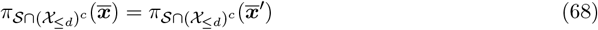

By assumption, 𝒳_*<d*_ ⊆ *S*, which implies that 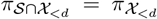, and thus (66) and (67) imply 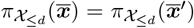, immediately followed by 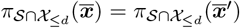. This, together with (68), shows that we have

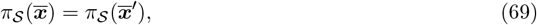

By assumption, 𝒮 is an indicator subset, so (69) implies 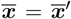. Since 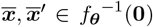 are two arbitrarily given points such that 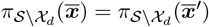, we hence see

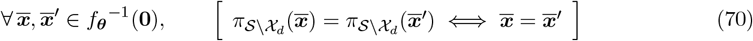

Now we have the proof. 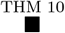

Now we shall start proving the main result Theorem 1. Collect all those blocks with sign-definite determinants in the diagonal of the structural decomposition form and put

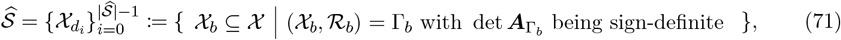

where 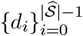 is a *strictly decreasing* subset of {0, 1, …, *B*}. Recursively define

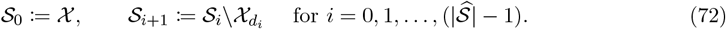

This forms a sequence of subsets 𝒮_0_ ⊇ 𝒮_1_ ⊇ · · · ⊇ 𝒮_|*ŝ*|_ such that

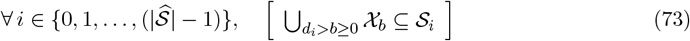

Obviously, 𝒮_0_ = 𝒳 (the set of all the chemicals) is an indicator subset, by definition. By Theorem 10, we inductively conclude that 𝒮_1_, 𝒮_2_, …, 𝒮_|*ŝ*|_ are also indicator subsets. Specifically, the smallest one is the one suggested in the main theorem,

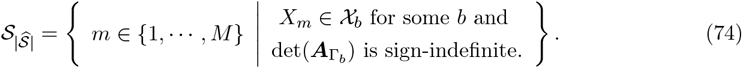

This completes the proof of the main theorem. 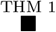

## D Appendix: The algorithm of identifying indicator species in a chemical reaction network

The framework of the algorithm is to repeatedly (1) find a buffering structure Γ with no emergent conserved quantities, (2) determine whether the corresponding det ***A***_Γ_ is sign-definite, and (3) reduce the network by eliminating Γ, until the network contains no non-trivial buffering structures with no emergent conserved quantities. This idea leads to an algorithm, and the pseudocode is given by **Algorithm 1** presented below.

### Algorithm 1: Algorithm of identifying indicator subsets

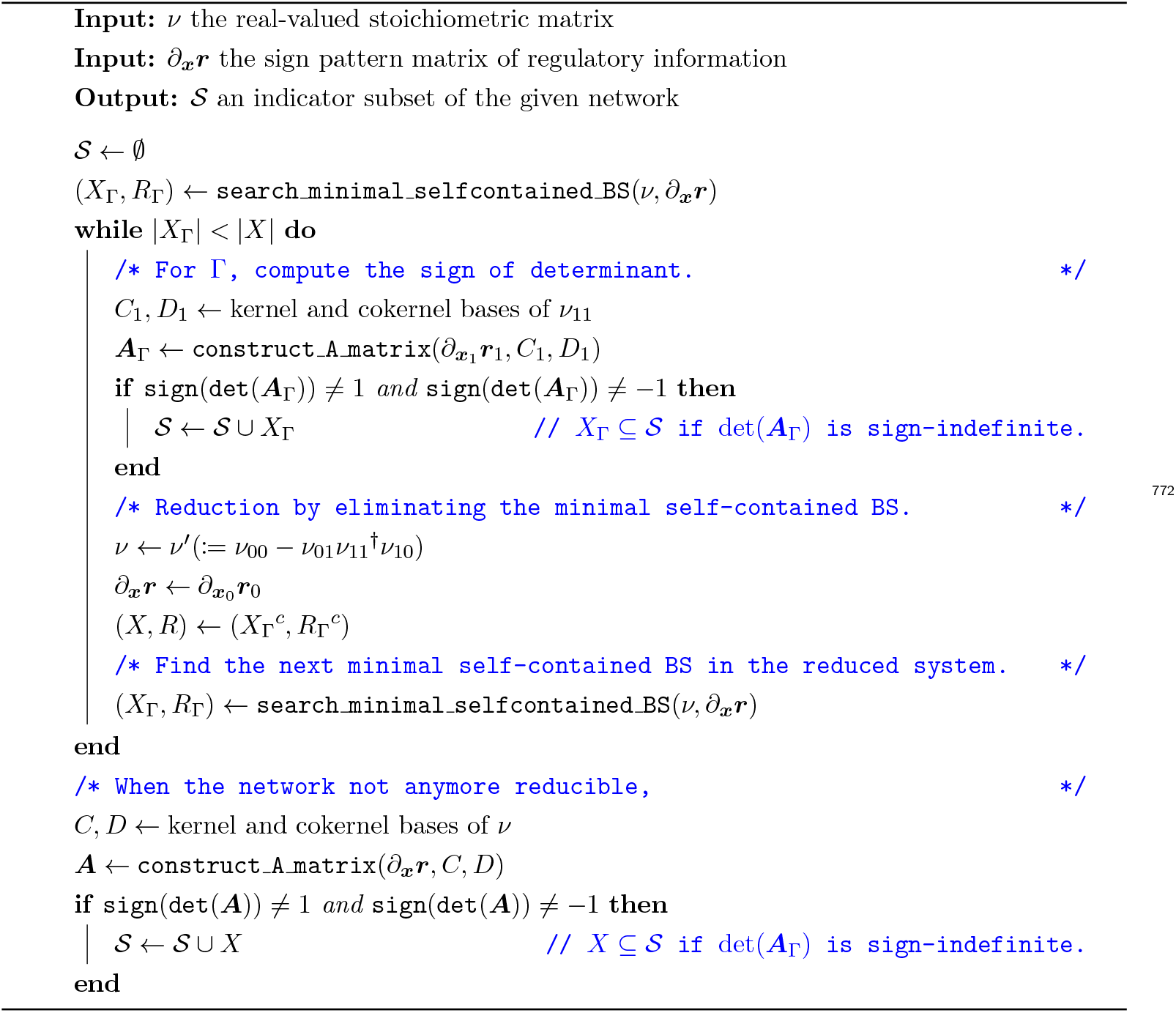

In `search_minimal_proper_BS`, the buffering structures are identified by following the strategy suggested by Yamauchi et al. [39] (with slight modification), and `search_minimal_proper_BS` chooses a/the minimal buffering structure with no emergent conserved quantities as the output. By *minimal*, we mean the number of chemical(s) (i.e., |*X*_Γ_|).

### Algorithm 2: search minimal selfcontained BS()

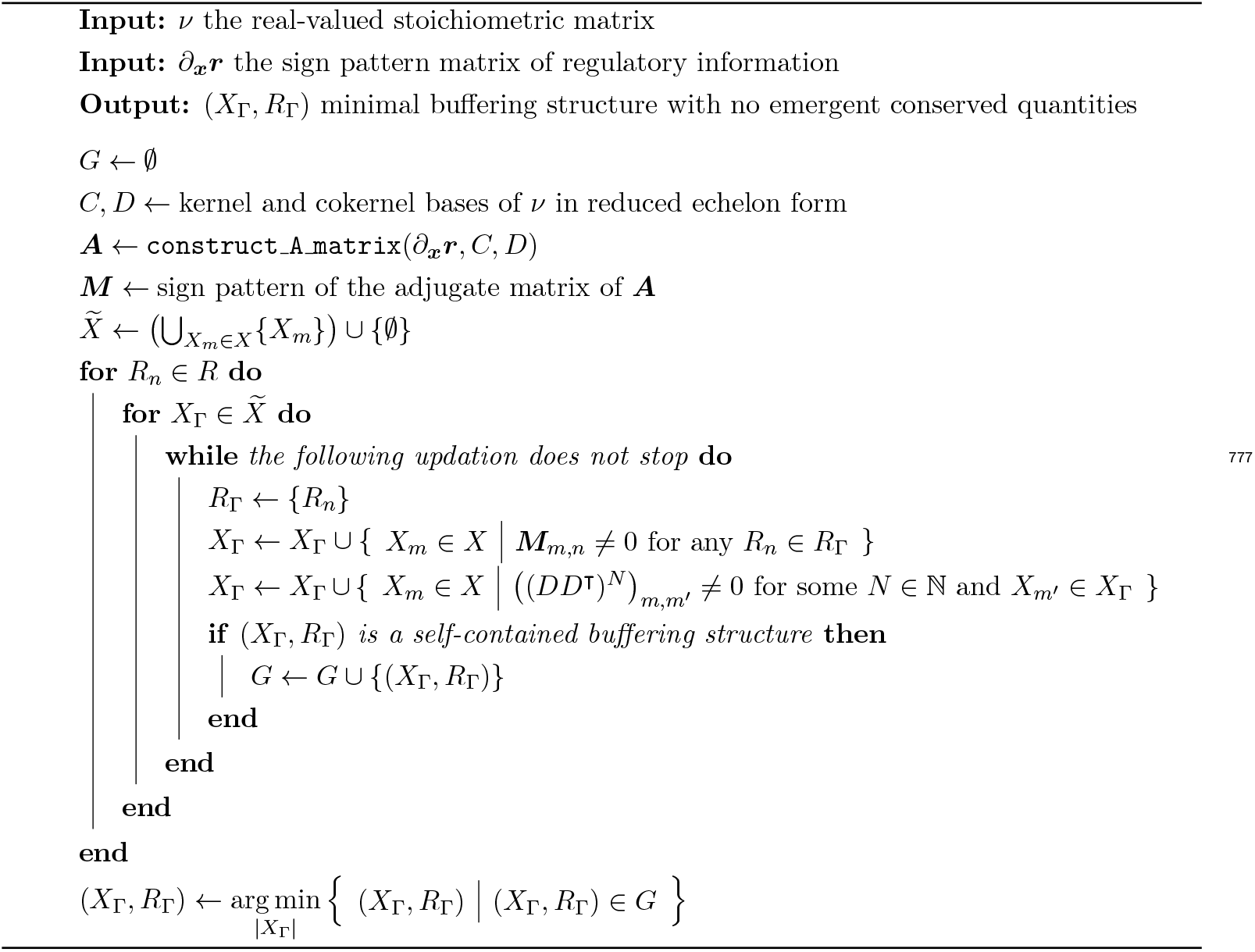

It is a problem both in **Algorithm 1** and in **Algorithm 2** that, given a matrix of each entries are in ℝ ∪ {+, −, ±} (i.e., fused with scall) how the sign of the determinant can be determined. This problem occurs in the evaluation of sign(det(***A***)) in **Algorithm 1**. This is also issued in **Algorithm 2**, when the sign pattern ***M*** of the adjugate matrix of ***A*** is being constructed.

For a square hybrid matrix ***K*** = (*K*_*ij*_) composed of scalars and sign patterns, we use a numerical strategy to determine the sign of its determinant. Roughly speaking, taking those *k*_*ij*_’s which are not scalars but sign patterns as variables, then det ***K*** is a smooth function. To check the sign, one can make use of optimization method to determine the maximum and minimum of arctan(det(***K***)). Note that arctan is considered so that the optimization procedure converges even when the determinant tends to ∞ or −∞. The sign of *k*_*ij*_’s are constrained in the procedure.

As an example for the clarification, for a hybrid matrix

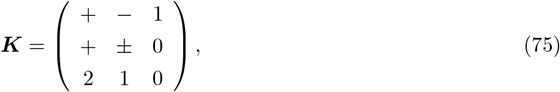

we consider a function

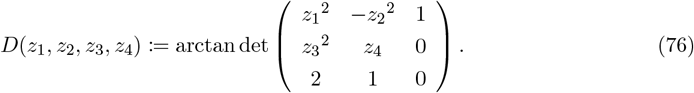

The exponentiation are given for the constraint on the sign patterns. Using an optimization method, one can evaluate 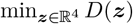 and 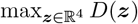, and sign(det ***K***) can thus be determined.

## E Appendix: numerical experiments on the running example

For readers interested in the details of the verifications of our theory by numerical experiments on the five-species running example network, more details are given in this section.

The distribution of phenotypic states are illustrated in Fig. 5. With the plot of PCA, it is clear that there are three clusters, which represent the three phenotypic states. Variation within the same phenotypic states are induced from the slight parameter variation. One can see that the overlapping of phenotypic states are severe under projection to some subsets even when the measurement noise is not present. Phenotypes 1 and 2 cannot be distinguished without using data of chemical A.

In addition, we evaluated the robustness of our method against measurement errors (Fig. 6). Recall that in the main text we estimated AMI and SILH scores from clustering results based on different subsets, fixing the measurement noise strength at *σ*^2^ = 0.15 and repeating the experiment 100 times (Fig. 4). Here, we performed a similar comparison between the results obtained using all chemicals (*A, B, C, D, E*) and those obtained using the predicted indicator subset (𝒮 = {*A, E*}), while varying the measurement noise strength *σ*^2^ from 0 (noise-free) to 1.5.

**Figure 5.**
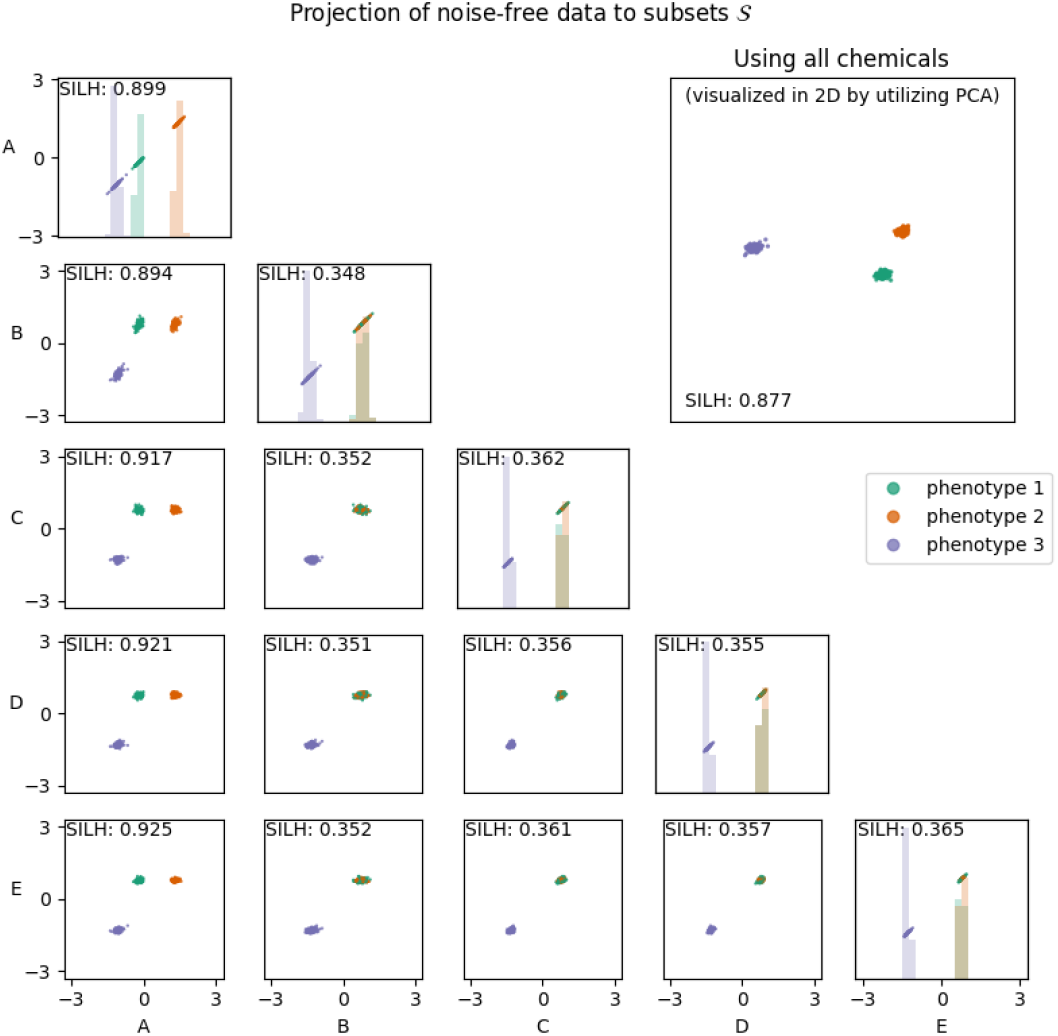
Visualization of the distribution of noise-free data, presented in a manner parallel to Fig. 3. The projection onto the first two principal components (derived from the noise-free data) clearly reveals three distinct clusters, which serve as the ground truth classification for subsequent experiments involving noisy observational data. Variations within each cluster reflect the differences in system states arising from parameter variability within the tissue.

**Figure 6.**
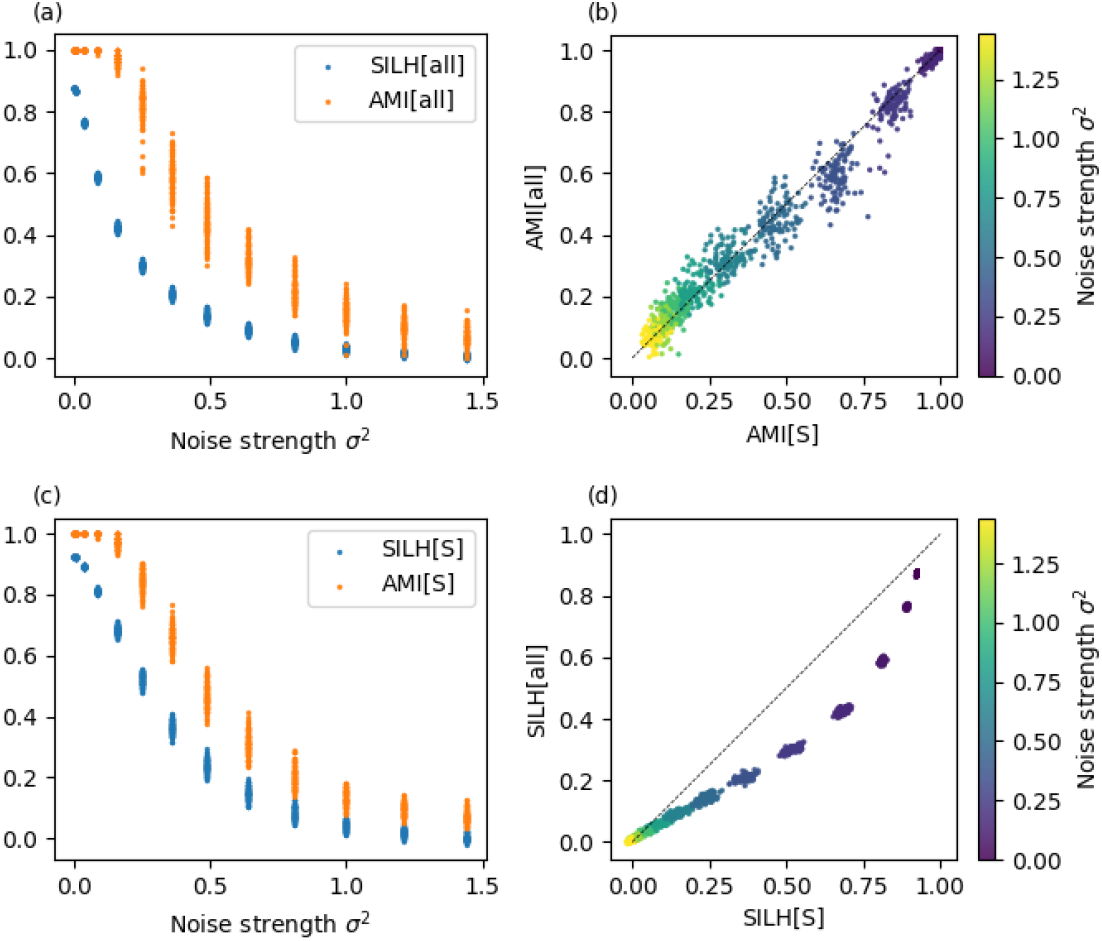
The test of robustness against measurement noises, comparing the performance of clustering when using all the chemicals and using the predicted indicator subset 𝒮 = {*A, E*}.

Both indices decreased as the noise level increased (Fig. 6 (a) and (c)); however, the rates of decline (i.e., ∂AMI*/*∂(*σ*^2^) and ∂SILH*/*∂(*σ*^2^)) were noticeably more moderate when only *A, E* were used, compared with using all chemicals. The robustness of employing 𝒮 rather than the full set is summarized in Fig. 6 (b) and (d). Specifically, panel (b) shows that classification based on 𝒮 achieves higher accuracy when the noise level is moderate (0 ≤ *σ*^2^ ≤ 0.75). At higher noise levels (*σ*^2^ *>* 0.75), the AMI obtained using all chemicals appears slightly higher than that obtained with 𝒮; however, the overall low absolute AMI values in this regime indicate that clustering has essentially lost discriminative power under such highly noisy conditions. On the other hand, for any noise strength *σ*^2^ setting, the SILH scores using 𝒮 stay higher then using all the chemicals. Namely, the distinguishability of phenotypic states is always higher if we only consider data of 𝒮.

